# An independent base composition of each rate class for improved likelihood-based phylogeny estimation; the 5rf model

**DOI:** 10.1101/2024.09.03.610719

**Authors:** Peter J. Waddell, Remco Bouckaert

## Abstract

The combination of a **t**ime **r**eversible Markov process with a “hidden” mixture of **g**amma distributed relative site rates plus **i**nvariant sites have become the most favoured options for likelihood and other probabilistic models of nucleotide evolution (e.g., tr4gi which approximates a gamma with four rate classes). However, these models assume a homogeneous and stationary distribution of nucleotide (character or base) frequencies. Here, we explore the potential benefits and pitfalls of allowing each rate category (rate class) of a 4gi mixture model to have its own base frequencies. This is achieved by starting each of the five rate classes, at the tree’s **r**oot, with its own free choice of nucleotide **f**requencies to create a 4gi5rf model or a 5rf model in shorthand.

We assess the practical identifiability of this approach with a BEAST 2 implementation, aiming to determine if it can accurately estimate credibility intervals and expected values for a wide range of plausible parameter values. Practical identifiability, as distinguished from mathematical identifiability, gauges the model’s ability to identify parameters in real-world scenarios, as opposed to theoretically with infinite data.

One of the most common types of phylogenetic data is mitochondrial DNA (mtDNA) protein coding sequence. It is often assumed current models analyse robustly such data and that higher likelihood/posterior probability models do better. However, this abstract shows that vertebrate mtDNA remains a very difficult type of data to fully model, and that dramatically higher likelihoods do not mean a model is measurably more accurate with respect to recovering key parameters of biological interest (e.g., monophyletic groups, their support and their ages). The 4gi5rf model considerably improves marginal likelihoods and seems to reverse some apparent errors exacerbated by the 4gi model, while introducing others. Problems appear to be linked to non-stationary DNA repair processes that alter the mutation/substitution spectra across lineages and time. We also show such problems are not unique to mtDNA and are encountered in analysing nuclear sequences. Non-stationarity of DNA repair processes mutation/substitution spectra thus pose an active challenge to obtaining reliable inferences of relationships and divergence times near the root of placental mammals, for example.

An open source implementation is available under the LGPL 3.0 license in the beastbooster package for BEAST 2, available from https://github.com/rbouckaert/beastbooster.

## Introduction

Nucleotide sequence evolutionary models typically assume that DNA mutations and substitutions are instantaneous stationary time-reversible Markov processes that occur at different rates across sites (Swofford et al. 1996, Felsenstein 2003). Some sites may be invariant/invariable, meaning that they cannot tolerate change. Other sites may change according to fixed relative rates, such as being collectively described by a gamma distribution. Mixture models can be used to avoid estimating individual site rates in order to calculate site pattern probabilities for a whole alignment of such sites. Such methods have become probably the most widely used likelihood, or site pattern probability-based, methods of estimating evolutionary trees (e.g., in PAUP*: Swofford 2000, BEAST: Suchard et al, 2018, BEAST 2.5: Bouckaert et al. 2019, RAxML: Stamatakis 2014, MrBayes: Huelsenbeck and Ronquist 2001). Here we look at the components of these models and then implement a novel extension.

Markov models of nucleotide substitution have undergone their own sequence of evolution since Jukes and Cantor (1969). Substitutions originate with and are proportional to the mutation spectrum in a neutral model of evolution. This assumes all types of substitutions are equally likely. Neyman (1971) and Felsenstein (1981) developed the basic mathematical machinery to apply such a process across a weighted tree to predict and fit all site patterns (e.g. by maximising the likelihood). An alternative machinery for certain classes of models, using generalised “distances” and ultimately fast Hadamard transforms, was developed in Hendy (1989) and Hendy and Penny (1993). The earliest assumption was a Poisson process of evolution (Jukes and Cantor 1969, abbreviated jc). An important step was allowing transitions (A:G and C:T changes) and transversions their own different rates (Kimura 1980, k2). Another generalisation was to mix k2 with unequal stationary base frequencies (Felsenstein 1981, 1993, Hasegawa et al. 1985, hky). Other combinations of particular sets of substitutions were explored and solved in closed form (e.g., giving the AG and CT transitions their own rates). However, since the **g**eneral **t**ime **r**eversible model (Lanave et al. 1984, Tavere 1986, Rodriguez et al. 1990, gtr) has desirable mathematical properties, and is easy to constrain to all its sub-models, it has become the common machinery for this whole class of models.

Another important evolutionary property is that a given site in a DNA sequence can have a different propensity to undergo a substitution. A set of completely invariant sites can be assumed and implicitly modelled (Hasegawa et al. 1985, Reeves 1992, indicated in shorthand as an i). Variable sites can have fixed relative rates chosen from a specific distribution. A gamma distribution is flexible and has desirable mathematical properties, but so too do other distributions such as an inverse-Gaussian distribution (Waddell et al. 1997). Such distributions of site rates were first modelled with distances (e.g., Jin and Nei 1990) and then to derive site pattern probabilities (Steel et al. 1993, Yang 1993, indicated in shorthand by g). Such approaches are called mixture models, as the final site pattern probabilities are a mixture of individual process that get integrated (summed up) to produce a single vector of site pattern probabilities. An alternative is to partition the sites into biologically meaningful sets, for example codon positions (Hasegawa and Kishino 1989, Shapiro et al. 2006) and combining the two is straightforward being implemented in many phylogenetic packages. Mixtures of invariant sites and gamma distributions were implemented independently using different machineries (e.g. Gu and Li 1996, Waddell 1995, Waddell and Penny 1996) and in the case of Waddell and Steel (1996) allowed the variable and invariant sites to have their own nucleotide frequencies.

A really significant and fundamental problem for phylogenies based on sequences is non-stationary evolution. The mutation process is driven mostly by a cocktail of DNA proof reading/repair enzymes, that is not homogeneous across time or genomic space, that is, the substitution process itself evolves. It is suspected of being perhaps the major outstanding issue for phylogenetic methods as total amounts of evolution increase (e.g., Lockhart et al. 1992a,b). It is often seen as a “long edges attract” problem, where two taxa independently evolve a distinct nucleotide composition, resulting in the underestimation of evolutionary distances due to increasingly matching character states not due to common ancestry (convergence). However, a potentially larger problem is “long edges repel”, for example, where two lineages evolve in different directions (Waddell 1995). In that case, application of potentially highly non-linear “distance” corrections (such as with a mixed invariant sites gamma distribution) can then result in massive relative overestimation of some path lengths; a particular problem in the anti-Felsenstein zone (Waddell 1995).

Here we implement into BEAST2 a model that starts with the machinery of the gtr4gi model, but then allows each of the five rate categories to have its own starting root frequencies (the gtr4gi5rf model). A common rate matrix made up of a stationary frequency vector and gtr rates of exchange is assumed for all variable sites, but, in future, this could be unlinked also. Via a simulation study, “practical identifiability” is assessed. This is the ability to return mean estimators and accurate credibility intervals across realistic parameter values and sequence lengths. Then, gtr4gi5rf is assessed on a range of real data. We consider how this extension of current models compares to other steps in the development of supposedly improved model of sequence evolution. Some specific results speak to how difficult it may remain to reliably infer, for example, clades of super-ordinal placental mammals, along with their divergence times, despite very long sequences.

A final short note on the added computational complexity of 4gi5rf models. Typically, with a 4gi model approximately 95% of the computational effort when recalculating likelihoods involves partial updates of the likelihood while about 5% involves the recalculation of the transition matrix. BEAST deals exclusively with rooted time trees so there is no added complexity there: ML implementations of this model would need to explicitly locate the root. The 4gi5rf model only adds a few extra multiplications at the root, so, in a binary tree of 2*n*-2 branches, this is just 2/(2*n*-2) (where *n* is the number of taxa). It also involves setting up 5 extra frequency vector priors, but with efficient proposals, a similar overall effective sample size (ESS) can, in theory, be achieved in a Bayesian implementation. We evaluate how well these extra parameters mix and whether chains need to be significantly longer to achieve similar effective samples sizes.

## Materials and Methods

The new model feature, which is the ability of each rate category to have its own independent root or starting nucleotide frequencies was implemented in the beastbooster package for BEAST2 (Bouckaert et al. 2019) and is accessed via the options in the supplementary BEAST2 input xml files. Amongst these options is use of the BICEPS package (Bouckaert 2022) to sample trees from the Yule-skyline tree prior based on a series of epochs, each with its own rate of lineage splitting (these are abbreviated “myule” for multiple Yule skyline, and give a high degree of flexibility of tree “shape”). Further, the “optimised relaxed clock” or ORC package was used to allow for unequal evolutionary rates amongst lineages (Drummond et al. 2006, Douglas et al. 2021). Another feature, very useful with shorter alignments where a particular type of substitution (e.g., A:T) might be rare or missing, is Bayesian Model Test or bmt (although Bayesian Model Averaging is a more accurate description). This enables reversible jumps into more or less parameterised models of DNA substitution helping to prevent over parameterisation (Bouckaert and Drummond 2017). A range of operators and types of chain were tried. It was found that MC3 did not seem to perform any better than multiple runs of MCMC with these data, so most analyses were based on four or more independent MCMC runs with convergence checked using Tracer 1.7.2 (Rambaut, et al. 2018) and DensiTree v2 (Bouckaert and Heled 2014). If there was convergence, but burnin was more than 10%, then burnin was estimated visually and input to LogCombiner (Bouckaert et al. 2019) to produce a concatenated chain file. Unless otherwise mentioned below, standard defaults proposals from BEAUti 2.7.5 seemed to work acceptably well.

Three biological data sets were examined in detail. The first “primate_mtDNA” is an alignment of 35 taxa and 1,832 protein coding mtDNA sites from four placental mammalian orders focusing on the phylogeny of Madagascar primates (Yang and Yoder 2004). The second “vertebrate_mtDNA” is an alignment of 44 taxa and 10,254 sites of the 12 heavy strand protein encoded mtDNA genes spanning major vertebrate lineages (Waddell et al. 1999a, 1999b). The third is “rag1gfib” or “ragfib_nuc” an alignment of 42 taxa and 1,032 nuclear encoded RAG1 and gamma fibrinogen nuclear encoded protein coding gene fragments sampled across all placental orders of mammals (Waddell and Shelly 2003).

In order to assess the potential impact of non-stationarity programs of the Interrogate package (Waddell et al. 2005) were used, in particular “FreqNuc.” This performs a set of tests of stationarity of base composition (e.g., the generalised least squares test of Tavere 1986, with grouping, SS5) and time-reversibility via the symmetry of pairwise divergence matrices (e.g., the likelihood ratio test of symmetry, G2sym). The full matrix of pairwise test results was visualised using a NeighborNet (Bryant and Moulton 2004) implemented in SpitsTree4 (Huson and Bryant 2006). These tests are performed on the pairwise divergence matrix of A to A, A to C, … T to T changes for each pair of taxa. The stationarity test evaluates differences in row versus column sums taking into account shared/correlated entries (leaving 3 degrees of freedom), while the symmetry test evaluates the equality of the six pairs A to C relative to C to A, etc (with six degrees of freedom).

To allow for independent nucleotide frequencies across rate classes, there are two ways this might be achieved. Since BEAST deals purely with rooted time trees, one way this can be achieved is by separately setting the root frequency parameters of that rate class. In the case of invariant sites, nothing changes down the tree and this root frequency is also the stationary (or equilibrium) frequency vector of these sites. However, for the variable sites, the overall result is a non-stationary non-time reversible model. To illustrate, let’s assume the frequency vector is nearly all A at the root, but the relative rate matrix on each edge of the tree is symmetric. This means the ultimate stationary distribution of this model will be an equal portion of A, C, G. T. If we were to visit each daughter node of the tree, ordered by the total amount of evolution to that point (that is, the sum of branch lengths along that path from the root), then what we would see is the expected base composition vector moving from all A to become increasingly uniform. The less the amount of change, the better the approximation to a corresponding time reversible model is expected to be.

The second way to allow each rate class to have its own base composition is to set a different frequency vector **f** for each rate class and make the root frequencies of that rate class the same. This model is a stationary mixture model. In this model, each time the frequency of any nucleotide changes, a new rate matrix needs to be exponentiated, and used to recalculate the corresponding transition matrices, and then recalculation of all likelihoods by multiplying the root frequency vector down the tree. This results in a substantial increase in computational cost. For the previous approach, unless the nucleotide parameter is in a stationary vector, only the set of multiplications down the tree is recalculated.

The acronym used to describe this model and its sub-models is a **t**ime **r**eversible Rate Matrix with **4**-equal sized site rate classes generated from a discretised **g**amma distribution plus a set of **i**nvariant sites, equals a tr4gi model. The new model adds 5 **r**oot **f**requency vectors, so we would write tr4gi+5rf or just tr4gi5rf. If also using BMT we would can add this abbreviation, e.g. bmt4gi5rf.

These multiple free root frequency models use pretty much the same prior distributions and proposals a standard stationary frequency vector would use. The gtr4gi5rf can initially be slower to converge simply because there are 15 more free parameters to sample. To achieve a similar sampling efficiency required proposal sizes and/or weights should be evaluated and adjusted as needed.

## Results

### Practical identifiability of the models and tuning

Supplement 0 shows the results of a well calibrated simulation study (Mendes et al. 2024), where we generate 100 random realisations of the hky4gi5rf model, then assessing how well the program was able to recover the parameter values used to generate each dataset. We used a fairly short sequence length of 500 to be conservative in our assessments. This assessment included how accurate the Highest Probability Density (HPD) 95% intervals were, that is, how often they contained the “true” generating value within their coverage. With 100 replicates and an expected value of 95, the marginal distribution should be a binomial with mean 95 and variance 100 × 0.95 × 0.05, thus a standard deviation of about 2.2 with a slight skew. Thus, 95% of coverage estimates in this simulation should fall from 91 to 99 with < 5% being outside this interval.

Overall, the coverage values presented in Supplement 0 were reasonable. Since there are 30 parameters (of which 24 are independent), we expect zero, one or two coverages to fall outside the 91 to 99 range (with this variable too having a marginal binomial distribution). We see that freqParameter 5.2 has coverage of 88 and freqParameter 2.4 has coverage of 90. A coverage as low as 88 is a bit rarely expected, but the fact that these are both part of symmetric compound priors (four value frequency vectors), where the other parameter coverages seem ok, gives some assurance that that specific parameter is not the issue.

One result noticed is the interplay of truly invariant = invariable sites, that is sites that absolutely cannot change, and the slowest evolving ¼ of the sites that can change. This latter fraction gets increasingly close to the four fixed site patterns of the invariant sites as the shape parameter of the gamma distribution goes to zero (so the shape of the gamma becomes a hyper-exponential curve). One manifestation of this is the often very strong correlation between the estimated proportion of invariant sites, pinv, and the shape parameter (Waddell 1995, Waddell et al. 1997). It is thus a problem for practical identifiability.

With the 4gi5rf model, these two rate classes can also assume a nucleotide frequency independent of each other. Because of this, pushing A high in the invariant sites can mostly be accommodated by dropping the A lower in the slowest evolving rate class, and so on. This can result in fairly broad marginal distributions of these parameters. This may contribute to poor mixing of the chain if proposals that adjust for these correlations are not made. Running a well calibrated simulation study assuming the 4gi model (not shown) we found that as long as the gamma shape parameter was greater than 0.1, we could usually untangle the proportion of invariant sites (consistent with Bouckaert 2020).

We also observed that the variances of frequency parameters, even in the same rate class, can be quite erratic (for example, one or two parameter variances much larger than the others). This was partially addressed by adjusting the weights on the proposals for these parameters and/or the proposal step size, and sometimes the adjustment compared to the standard model is large (∼ 50 to 100 times higher weights on some frequency interchange proposals were helpful in the real examples below).

The priors for the calibration simulations were chosen to be quite broad. However, for the mtDNA data mentioned below, we occasionally see at least one of the parameters outside of the simulation range. In particular, estimated frequency of G in the gtr4gi model might come in near the extreme of its simulated range (e.g., 0.06). However, the corresponding parameter of the gtr5gi5rf model might drop to 0.02 and the sampling distributions of the root frequency parameters might also fall into the extremes. These seem to be related to interaction of the model with the non-stationary nature of the data, as discussed below.

### Application to primate_mtDNA

Figure 1 shows the marginal distributions of the rate and frequency vectors of the new 4gi5rf model run on this data. It was found that mixing of the model can take a bit of tuning to achieve acceptable sample sizes within 10 million steps, with weight adjustments helping considerably. After weight adjustment there was still an association between ESS and the marginal variance of a parameter, despite the proposal magnitude being automatically tuned. This issue may arise when the overall vector of proposals is optimised by scaling, but not the individual elements. That appears to be the case as the standard deviation of individual elements of the frequency vectors was observed to vary by over a factor of four. Another interesting pattern is how the root frequencies of the two slowest rate classes (pinv and gamma rate category 1) mirror each other.

**Figure 1.**
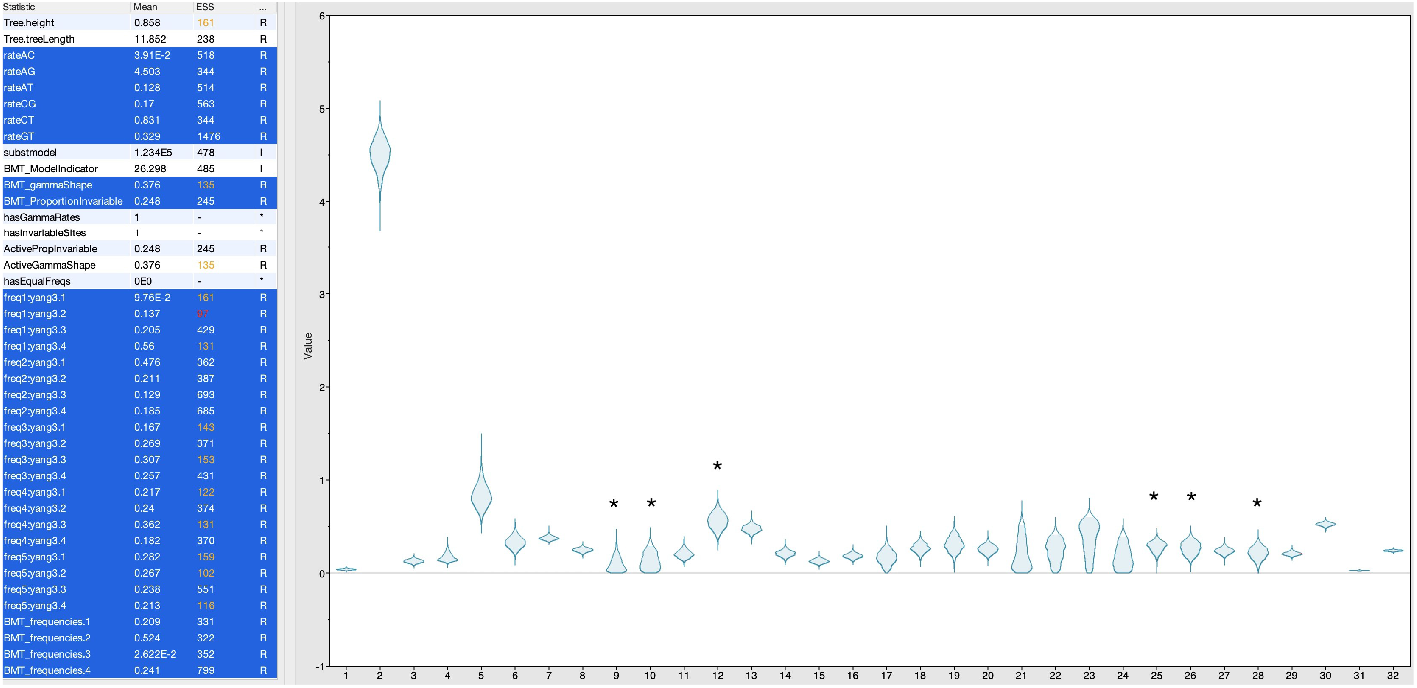
The marginal distributions of the gtr4gi5rf model applied to primate_mtDNA. Left to right, the six relative mutation rates, the gamma shape then pinv, then the five vectors of root frequencies (slowest to fastest rate class), and, finally, the stationary frequencies. With the frequency parameter names, the last digit indicates the nucleotide, A=1, C=2, G=3 and T=4. A numeral in the first part of the name of a frequency vector (for root frequency vectors only), indicates the rate class, with invariant sites = 1, slowest quartile of a gamma distribution = 2, etc. The asterisks indicate examples of root frequency parameters with much larger variances than their neighbours, often leading to their having lower ESS estimates. Note also the extremely low frequency of G in the stationary frequency vector and the inverse relationships of frequencies in the first two root frequency vectors.

In this example, many of the variances of the root frequencies for the rate classes are large and those of the stationary frequencies sharply different and precise (very clear for G).

Another issue that seems to get worse with more rate class frequency vectors and shorter alignments is over-parameterisation of the symmetric rate matrix. It is tempting to automatically use gtr, assuming that the worst that will occur is a generally higher variance of parameter estimates. However, if one of the rate matrix entries has a very low value, then its estimation might be largely based on a site pattern occurring just once in the data, so its reconstruction might be quite dissimilar on different trees. In this case strongly bimodal marginal distributions of the rate parameter can arise and mixing can be poor. A potential solution is to apply bmt to the free parameters namely, the rate matrix entries, stationary frequency vector, pinv, gamma shape, etc.). It may be useful to apply bmt also to all the new frequency vectors, but here that was not necessary to address the problems mentioned. Application of gtr4gi5rf leads to an increase of the lnL relative to the standard gtr4gi model by about 335 lnL (natural log) units for an extra 15 free parameters, which is considerable. Despite limited data, the posterior was also superior by about 144 lnL units and completely separated from that of the standard model. While the full priors have not been checked for their integral summing to 1 (that is “normalized”), the priors of the extra 5rf parameters (which are Dirichlet distributed) are normalized. Therefore, it seems safe to compare at least this pair of nested models posterior distributions.

### Does this model really make much of a difference?

A comment on the new model, perhaps inspired by an early misapplication, was to the effect that the model dramatically changes likelihood, but does not change the inferred trees much, which makes the new model not particularly interesting. This is a potentially valid point which may be objectively addressed by looking in detail at the impact this model to the tree, that is topology, support values and node heights, relative to other proposed models that have been judged important advances over time. Notable steps in the evolution of likelihood models include that of Jukes and Cantor from 1969 (the Poisson or jc model), Kimura 1980 (transitions versus transitions, k2), Hasegawa et al.1985 (adding unequal base frequencies, hky), general time reversible model (further splitting transitions and transversions into six relative rates, gtr), the gtri model (adding an invariant sites rate class), the gtr4g model (a four category gamma site rate model), and, finally, the gtr4gi model with variable sites following a gamma distribution plus a portion of invariant sites.

Figure 2 shows the relative likelihoods and posteriors of our seven model steps applied to the primate mtDNA data. From left to right the models fall in the same order as the previous paragraph with 4gi5rf at the far right. In this lineage of increasingly complex models, the new gtr4gi5rf model shows an appreciable likelihood improvement, comparable that seen with most previous model expansions, but nowhere as large as the jump induced by adding invariant sites alone, which stands out as the largest single improvement.

**Figure 2.**
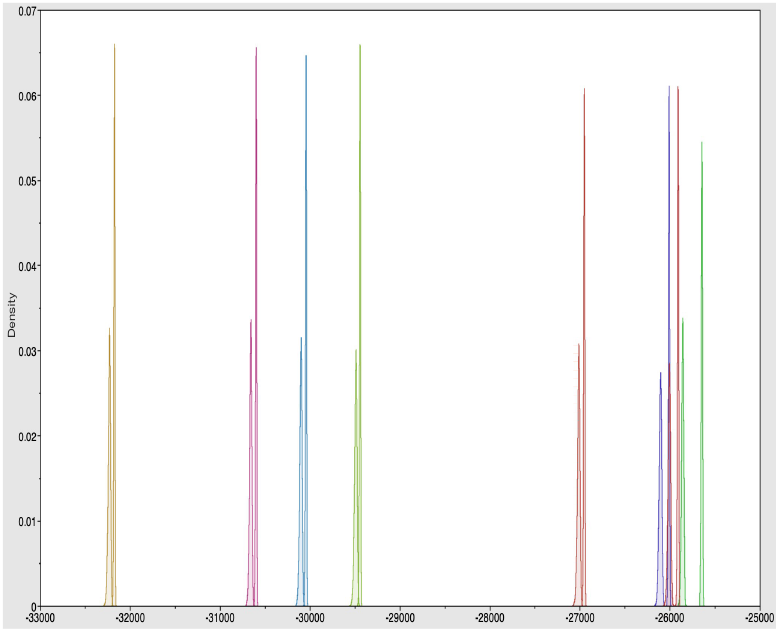
The relative likelihoods (higher peaks) and unnormalized posteriors (lower peaks) of seven likelihood models applied to the primate mtDNA with orc and myule priors. From left to right they are jc, k2, hky, gtr, gtri, gtr4g, gtr4gi and gtr4gi5rf models. The *x*-axis is in lnL units. Note, the usual caution necessary when interpreting comparisons with unnormalized parameter priors, yet it is clear the new model adding 15 parameters, improves the likelihood considerably. In addition, the comparison of the posteriors of some models, such as the 4gi model to the 4gi5rf seems reliable, since the former is a sub-model and they differ by 15 free frequency parameters which have normalized priors.

Before discussing what these improved likelihood and posterior probability models might add to our biological understanding, it is important to justify our expectations of what the “true” mtDNA tree is likely to be. The focus is on those parts of the phylogeny that alter substantially between models. This is only recently possible by combining the results of several large nuclear genomic studies, along with a simple but robust statistical test of the likely species tree (Waddell et al., 2002). Firstly, the root is almost certainly between Primates and the other three outgroup orders. Secondly, it is still difficult to be certain which resolution of the outgroup orders Carnivora, Perissodactyla, and Cetungulata is correct, not least since even genome scale sequence data can suffers from issues of non-stationarity and these clades are interrelated by short internal branches (Waddell et al. 1999b, 2001, Springer et al. 2001). The expectation that orders within Laurasiatheria were involved in a rapid diversification near the KT boundary is supported by the lack of resolving Line/Sine insertions amongst these clades (Kreigs et al. 2006, Nishihara et al. 2006). While Nishihara et al. suggest the correct arrangement is (Perissodactyla, Carnivora) we remain cautious due to ascertainment bias issues with some such characters (Waddell et al. 2003). Within Primates the consensus is that *Daubentonia*/aye aye is indeed the first branch of the endemic Madagascar primate clade (so closer to lemur than *Loris*+*Galago* are), something corroborated recently by Marciniak et al. (2021). The exact location of *Lepilemur* on the tree had been a bit ambiguous, but Marciniak et al. (2021) resolve it as ((((*Microcebus,Lepilemur*):92,*Propithecus*):60,*Eulemur*):98,*Daubentonia*), where the number is the percentage of many thousands of gene trees supporting each resolution. As long as these gene trees are inferred in a non-biased way, this in turn yields highly significant support for all nodes assessed by the three-taxon rooted multinomial coalescent species tree test of Waddell et al. (2002). For the resolution within the lemur clade there is an ALU insertion phylogeny in McLain et al. (2012) that supports the tree (((*Lemur, Hapalemur*), *Eulemur*), *Varecia*) with >95% support for each node, as assessed by the three taxon test of Waddell et al. (2002). Merging these results, the genera level subtree is (((((*Microcebus* + allies, *Lepilemur*), *Propithecus*), (((*Lemur, Hapalemur*), *Eulemur*), *Varecia*), *Daubentonia*), (*Loris, Galago*)).

In terms of the age of clades, the fossil evidence overall suggests, with reasonable confidence, that nearly all or nearly all orders of placental mammals arose 55 to 65 mya (millions of years ago). To date there also remains a complete lack of any definite placental mammal fossil older than 70 mya, suggesting placental superorders did not arise deep into the Mesozoic. Ages around 100mya, which coincide with events like the opening of the South Atlantic, have become widely accepted (Waddell et al. 1999b,c, with corroboration in Waddell et al. 2001 and Springer et al. 2001, for example). However, Kitazoe et al. (2007) show that the use of apparently valid alternative analytic methods reduce the inferred root age of placentals from whole mtDNA from ∼127 mya to ∼84 mya.

Using DensiTree2 to compare sampled tree-sets, a full set of comparisons in our series of models are presented in Supplement 1. If a change is most likely to be an improvement, it will be marked with an (+) and if likely to be a deterioration (-) in the text to follow. If it is unclear or is neither a correction or error it is marked (?). Figure S1.1 shows the jc and k2 tree-sets are quite similar, and node heights are nearly collinear There are two clades that have altered support sufficiently to alter the clades of the mcc tree. The bear (*Ursus*) is no longer sister to the cat (*Felis*) and is now strongly sister to the dog (*Canis*) (+), and *Lepilemur* has moved sister to (*Microcebus, Mirza, Chierogaleus*) (+). In terms of support that does not change the mcc tree, three clades that appear to be correct have all moved to much higher posterior support values (+). With k2 versus hky, very little changes; something of a surprise given the likelihood improvement and the fact that the base composition of mtDNA is fairly extreme (S1.2). Going from the hky to gtr the relative node ages deeper in the tree are decreasing noticeably (S1.3). There are also quite a few clade changes, with nearly all of them problematic. The hky to gtr mcc tree flips from strong support for *Daubentonia* sister to all other Madagascar primates to substantial support (pp∼0.8) for it deeper in the tree (-). Other problematic changes include more support for a non-monophyletic Primates (-), and, perhaps, far too much confidence in the subtree ((Cetungulata, e.g. Bos, + Carnivora, e.g. Canis), Perissodactyla, e.g. Equus) (?).

The next step, adding invariant sites for the gtri model sees a truly massive boost in likelihood. There is also a large (∼40%) overall increase in branch lengths, with those deeper in the tree increasing the most (S1.4). There are also four marked changes in support for clades. The mcc tree regains *Daubentonia* in the correct position (+) with considerable support (pp ∼0.8). Also appearing is a shift in the root. Unfortunately, this is incorrect, but gets substantial support (pp ∼0.75). Another increase in support is for *Lepilemur* in the correct position (pp ∼0.55 to 0.85) (+), and the correct monophyly of the Strepsirrhines is reasserted (pp ∼0.6 to 0.9) (+). The change from gtri to gtr4g sees a further increase in likelihood and a massive increase in edge lengths (by a factor of nearly 4.5). There is also a considerable increase in the relative age of the deeper nodes, indeed, the Jurassic Monkey Hypothesis is literally accurate. That is, using the reliable horse-rhino calibration of 55mya (Waddell et al. 1999b), the inferred age of the anthropoids (monkeys) in this tree is about 145.0 mya, which is essentially on the undefined boundary of the Jurassic with the Cretaceous. The gtr4gi model results in a small difference to the gtr4g model (S1.6). The only notable change is a decrease in support for the arrangement (*Eulemur*, (*Hapalemur, Lemur*)); apparently a correct clade (-).

Finally, we come to the comparison of the new gtr4gi5rf model to the incumbent gtr4gi model (figure 3). Here it is clear that there is a sharp decrease in the relative ages of the root and the monkeys to more derived clades. There are also six substantial changes in clade support, of which five change the mcc tree. The first is that *Daubentonia* goes back into its correct location (+) with substantial pp (from ∼0.05 to 0.7) (+). The second is that the root position in the mcc trees is, for the first time, correct and shows a monophyletic Primates (pp ∼0.00 to 0.5) (+). The third is that the association of Perissodactyla with Carnivora switches to that of Perissodactyla with Cetungulata (Euungulata) as pp goes from ∼0.3 to 0.9 (?), but most readily supported by minimizing evolutionary ecological shifts). The fourth is that *Lepilemur* moves away from *Microcebus* and closer to *Lemur* with pp ∼0.1 to 0.6 (-). The fifth is an increase in the support for the anthropoids (monkeys/apes) from ∼ 0.7 to 1.0 (+). The last is that the correct clade (*Lemur, Hapalemur, Eulemur*) with pp ∼0.6 going to 0.45 is replaced with (*Lemur, Hapalemur, Varecia*) pp ∼ 0.35 to 0.5 (-).

**Figure 3.**
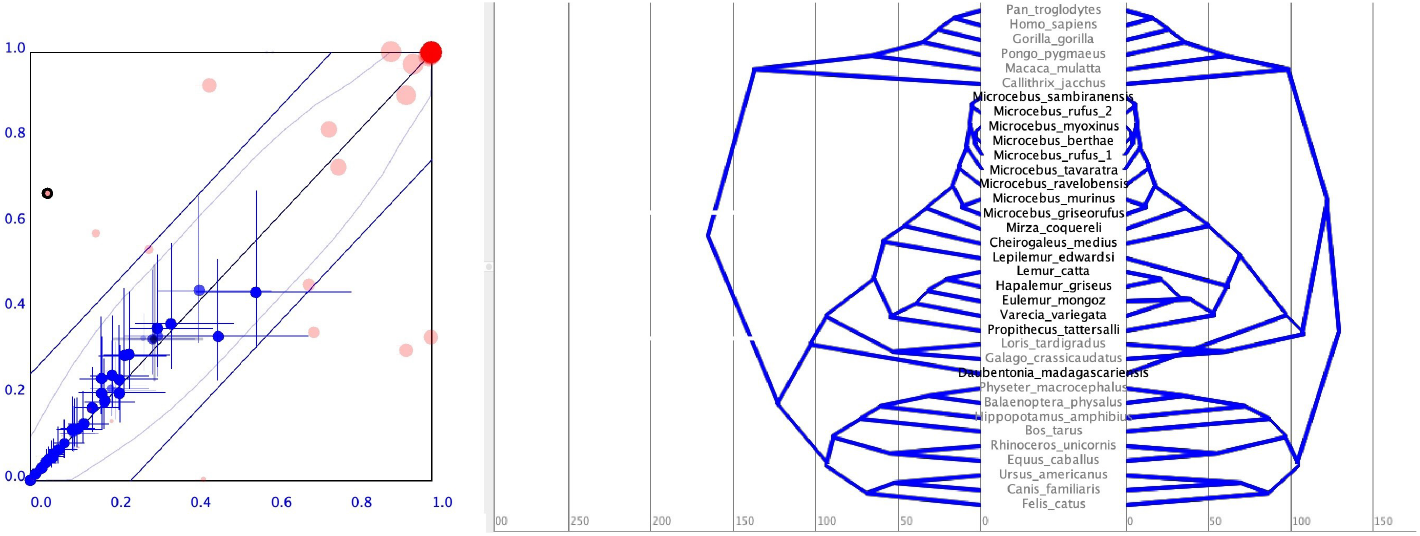
(a) A pairwise plot of clade support values (pink) and divergence time estimates (blue crosses representing 95% coverage intervals). The lower likelihood model (gtr4gi) appears on the *x*-axis, and the highest likelihood model (gtr4gi5rf) appears on the *y*-axis. (b) A mirrored mcc tree plot with gtr4gi on the left and gtr4gi5rf on the right. Here the scale is implied millions of years ago (mya) based on a single horse-rhino (*Equus-Rhinoceros*) calibration of 55 mya. Support for the monophyly of primates with the gtr4gi5rf model is ∼0.5. In contrast the gtr4gi based model is 100% convinced primates are not monophyletic.

Estimated node ages deep in the mcc trees are again calibrated via the horse/rhino split at (55mya). This yields root ages of 118.0:jc, 115.5:k2, 115.6:hky, 105.8:gtr, 118.2:gtri, 167.6:gtr4g, 170.3:gtr4gi and 128.6:gtr4gi5rf mya, respectively. The date that seems most unbelievable is the 170 mya for the gtr4gi model. This model suggests near humans (monkeys) walking with non-avian dinosaurs. While this is a favourite of Noahphiles, and might even be expected reported on Fox News, for example, to put it mildly, this contradicts considerable scientific evidence (-). By contrast, there are 170 minus 128.6 = 41.4 million good reasons why the new 4gi5rf model results are an improvement on those of the incumbent model 4gi. Interestingly, the biologically most realistic age for the placental root (∼105.8 mya) is produced by the gtr model. Usually, the more complicated models will infer more changes deeper in the tree (e.g., Waddell and Steel 1996); later we will discuss why this expectation can be incorrect in some situations.

Note, none of these trees resolved the position of *Propithecus* in concordance with the nuclear genomic results (i.e. closer to *Microcebus* than to *Lemur*). This brings us to a potentially uncomfortable point; which tree do other popular methods of phylogenetic estimation infer. Compared to the gtr4gi tree, unweighted parsimony favours Euungulata (?), and puts *Propithecus* even closer to the true lemurs than *Varecia* (-). NJ and BME with Hamming distances, a favourite method to resolve short edges affected by coalescent effects with low divergence (Waddell et al. 2011), again compared to 4gi places *Daubentonia* correctly (+) but also places cat sister to bear (-). In contrast, BME with LogDet distances, did not do particularly well. *Daubentonia* is placed deeper (-), cat and bear as still sister (-), *Lepilemur* moves 1 internode deeper in the tree (-), *Propithecus* and *Varecia* interchange (-). Increasing pinv in steps of 0.1 until negative edge lengths are encountered did not result in a better tree, indeed at pinv = 0.4 (pinv = 0.5 gave infinite distances), *Hippopotamus* moves one internode deeper (-), and the perissodactyls move sister to Carnivora (?). Using gtr distances with invariant sites = 0.326 and shape = 0.499 (their mean values from the gtr4gi BEAST model) results in an even worse tree with *Hippopotamus* moving another step deeper in the tree (-). Overall, there is nothing here to suggest that alternative methods resolve the position of *Propithecus* in agreement with the nuclear consensus, leaving this result potentially due to biological factors such as a failure of the mtDNA to coalesce in accordance with the species tree, lateral transfer of the mDNA, or a failure of essentially all phylogenetic methods to recover its correct position.

### Why does the gtr4gi5rf model make such pronounced differences?

It might be assumed that the primate_mtDNA data is fairly well behaved, as primates seem correctly located in whole mtDNA phylogenies (e.g., Waddell et al. 1999b). In fact, while the amount of evolution is nearly an order of magnitude less than across all vertebrates, the direction and extent in which some of the taxa (e.g. *Microcebus* versus *Pongo*) have shifted mutation spectrum seems similar (see figure 4 and S2.8.). Thus the rate at which some of these taxa are shifting away from each other would seem much faster than across most vertebrate mtDNA. It is known that estimating reliable trees from whole mtDNA at that level can be very challenging (e.g., Waddell et al. 1999a, 2001). The new model seems to be doing well for some of the deeper parts of the tree by adjusting when a lineage goes in a strongly different direction, in terms of base composition, to that in nearby branches. The monkeys do this, and another example is *Daubentonia*. Indeed, when the fastest evolving sites are considered (and they constitute most of what defines the relative likelihood of these trees), their root composition is fairly bland, but the stationary composition is hyper-monkey in its direction. This might be why, further away from the root, mistakes such as the location of *Lepilemur* appear.

**Figure 4.**
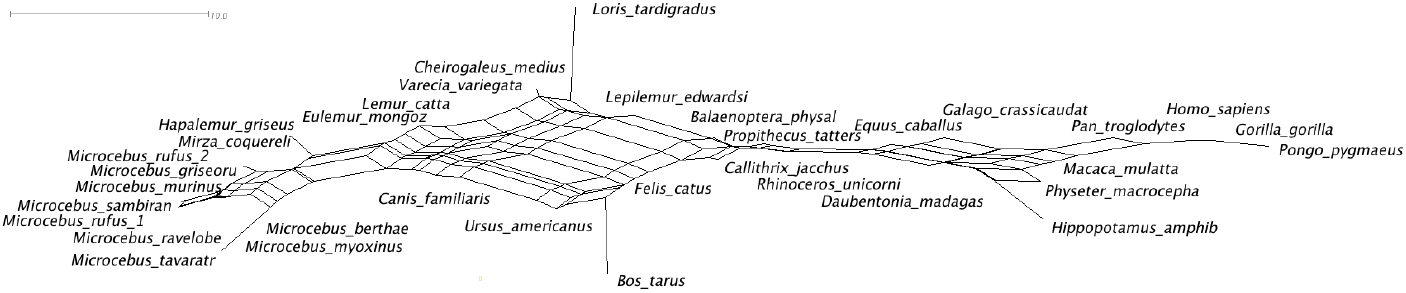
A NeighborNet visualisation of the pairwise distances induced by a SS5 pairwise test of base-composition stationarity. The percentages of A, C, G and T for *Microcebus rufus* 1 are 31 25 14 31, while those of *Pongo* were 29 34 13 25. Additionally, those of *Loris* are 33 27 12 27, and those of *Bos* are 32 28 14 26. The major left right trend is due mostly to increasing C and decreasing T, while that from *Loris* to *Bos* is mostly due to G increasing. When fitting the gtr4gi5rf model to these sequences, the stationary composition was 20 53 3 24, while the root composition of the fastest rate class was 29 27 24 20. The stationary composition is well beyond the observed composition of any of these taxa, but in the direction that the anthropoids (monkeys) are headed relative to *Microcebus*. In contrast the gtr4gi model has stationary frequencies 36 37 6 21. More such plots for each data set are shown in supplement 2. The scale bar indicates 10 units apart. Since the test statistic converges to chi-square with 3degrees of freedom, about 7.8 units apart is significant at the 95% level.

Another interesting feature of the data is that it appears most of the change in evolutionary direction of the taxa in primate_mtDNA can be explained by base composition shifts alone as tests of symmetry give very similar results (compare figure S2.1, S2.2). This suggests the gtr model might remain a reasonable approximation to the process of evolution, albeit with changes in the base composition vectors. Further, this association is closest at third position sites, still very clear at first position sites, and least clear at second position sites (which make the least contribution to the relative likelihood of trees), see Figure S2.3-S2.5. The pattern seen with the second position sites suggests that monkeys, including the readers, are being driven into somewhat radical amino acid changes by a mutation frequency spectrum that came into play with the evolution of the group.

Fitting the 4gi5rf model to first, second and third codon position sites separately further illuminates what is occurring The relative mean marginal log likelihood of the bmt4gi and bmt4gi5rf models at the first position sites were 6139.2 vs 6076.9 (for a difference of 62.3). At the first positions, the free root frequencies show the same patterns mentioned before, generally large variances with the invariant sites showing a frequency profile close to the reverse of the slowest quartile of the variable sites (here 17 18 27 37 vs 51 18 16 14).

At the third position sites, mixing was slower than desired (required burnin varied from 1 to 3 million steps) and one of six chains failed to converge. At these sites the inverse relationship of invariant sites (proportion only ∼0.015) and the slowest quartile was markedly less pronounced and frequencies across rate classes were more heterogeneous.

Partition by codon position improves the log likelihoods and posteriors by over 1000lnL units compared to the unpartitioned mixture models used earlier. The differences of the 4gi and 4gi5rf models are reduced when partitioned by codons, and, compared to the all-sites 4gi5rf model neither of these codon position models improve the biological assessment of the tree. Both codon 4gi and 4gi5rf models mis-root the tree with monkeys sister to all others. In terms of strongly different clades, there are just two. The codon-4gi5rf model strongly (∼0.85) favours the correct position for Daubentonia (+), as the codon-4gi model just as strongly favours an incorrect position (∼0.85). That situation is nearly exactly reversed with *Lepilemur* as seen previously in the unpartitioned mixture models, so a (-) for the codon 4gi+5rf model. Despite the strong increase in likelihood, character partitioning has not improved biologically fundamental phylogenetic parameter estimates.

### Adding in time and clade constraints

Adding in constraints on both clades and time can change how models behave. The current data is often run with constraints in place. It is not clear if this is to avoid the problematic “Jurassic Monkey” result. The constraints added in Yang and Yoda (2003) are not all directly based on easily interpretable fossil data with some an amalgam of commonly held expectations/beliefs. For example, their constraints on the age of Primates; the fossil record does not directly exclude markedly older ages (suggesting a fat tail to older ages), whereas for the age estimate of *Perissodactyla* the fossil record addresses this more directly, conforming more to a normal distribution in expectation (Waddell et al. 1999b). Infinitely strong constraints force the model into solutions, even if the model equally strongly prefers another clade. This can lead to a very different final result when comparing models. The constraints imposed by Yang and Yoda are not just on the age of nodes, but also on the monophyly of clades.

Figure 5 shows results for the fully constrained gtr4gi versus gtr4gi5rf models. A large increase in the log likelihood remains at about 340 ln units. The total tree heights are much more similar, but there is still a moderate effect at the root (which is being particularly influenced by the constraints on Primates). Here, the tree itself is similar to the unconstrained 4gi5rf tree, with circles showing three local alternatives encountered earlier.

**Figure 5.**
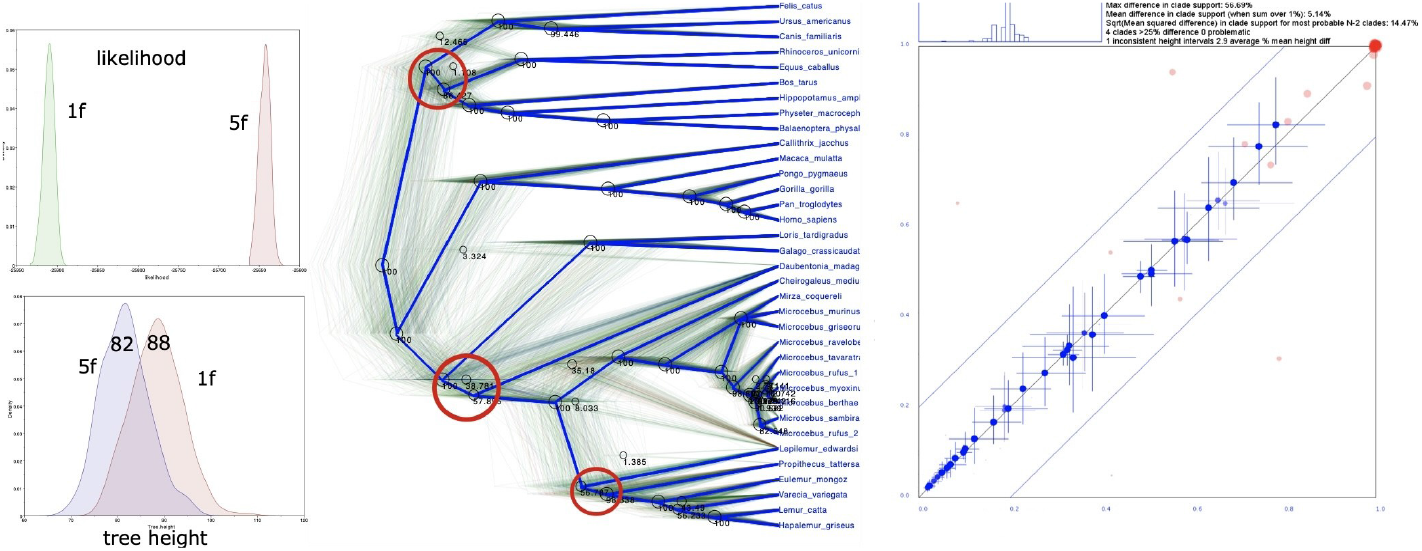
A comparison of the gtr4gi and gtr4gi5rf models applied to primate_mtDNA with multiple constraints in place. (a, top left). The distributions of the marginal likelihoods of the two models, (b, bottom left) Distributions of the tree heights of the two models showing their means (c, middle) The mcc tree for the gtr4gi5f model, superimposed on the DensiTree plot of all sampled trees, and, (d, right), a plot of the clade support values (posterior proportions, red) and the 95% credibility intervals of the node heights of the two models (see https://www.beast2.org/2020/04/20/comparing-tree-sets.html for a description of how this plot was generated).

### Application to a really hard phylogenetic problem, vertebrate_mtDNA

A good example of a really hard to analyse data set is the alignment of concatenated mtDNA protein coding sequences from Waddell et al. (1999b,c). The data span the major lineages of vertebrates. Quite profound problems in recovering correct clades occur within mammals, birds, and deeper in the tree with various fish. The gtr4gi5rf model chains converged relatively quickly (less than 1 million steps). Run out to 10 million steps the chain achieved a posterior ESS of 354 or more in all cases. Combined, the posterior ESS was over 1700. There was one sampled parameter with an ESS of 115 (AG), while three of the stationary frequency parameters were 83 to 187. For the 4gi model, one of the chains got stuck and was replaced. The improvement in the likelihood for the 5rf (15 extra free parameters) was ∼1870 lnL units. The stationary frequency of G with the gtr4gi5rf model was again very low at ∼0.03 (see figure 6). The slowest 2 rate class vectors showed strong inverse relationships. The frequency of A in the fourth rate class was surprisingly high at 0.913.

**Figure 6.**
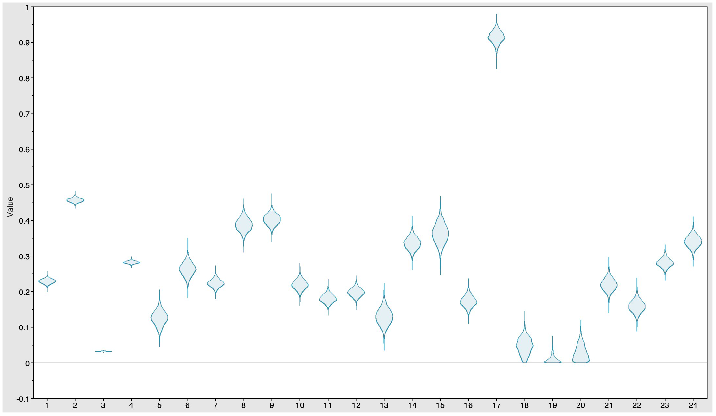
Violin plot of the frequency parameters for the gtr4gi5rf model applied to vertebrate_mtDNA. From the left, the first four parameters are the stationary frequencies of A, C, G, and T, then A of the first rate class (invariant sites), C of the first rate class, and so on.

**Figure 7.**
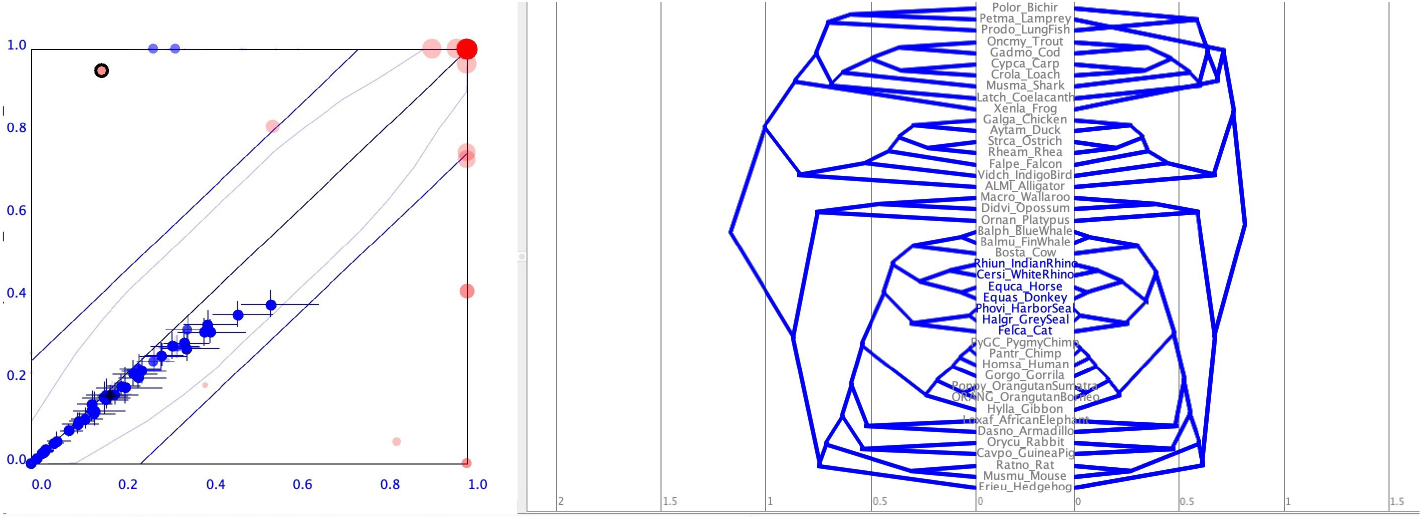
(a) A pairwise plot of clade support values (pink) and divergence time estimates (blue crosses) for trees based on vertebrate_mtDNA. The lower likelihood modelgtr4gi appears on the *x*-axis, and the higher likelihood model gtr4gi5rf appears on the *y*-axis. (b) A mirrored mcc tree plot with gtr4gi on the left and gtr4gi5rf on the right.

Unfortunately, the gtr4gi5rf model, while it strongly increases the likelihood of the data, does not result in any more biologically correct clades appearing in the mcc tree (the current biological understanding of the true tree is shown in figure S2.8_2). Here the model is again compressing the relative divergences near the root of this tree. In this case this is not consistent with the fossil record and may be due to parts of the data (such as third position four-fold degenerate sites) close to saturation. The new model does do one thing that might point to more promising results for this data with improved models of non-stationary base composition; it shifts the Lamprey deeper in the tree, the true biological outgroup to all the other taxa.

A look at the visualisations of pairwise SS5 stationarity and G2sym symmetry tests shows that these taxa definitely have issues that seem to stem from non-stationarity. By comparing the locations of overlapping taxa in the primate_mtDNA data with those in vertebrate_mtDNA, it is seen that the taxa in the primate data span most of the extremes of the vertebrate data. However, it is likely to be the combination of non-stationarity and a higher level of “saturation”, particularly in transitions, that makes the true phylogeny unrecoverable with present models. The term “saturation” is used too liberally in many publications. A more quantitative look at “saturation” is undertaken in Waddell (1995) where signal-to-noise ratio is compared under the model. It is shown, that this critical parameter, which translates directly to the recoverability of trees with methods such as distances and generalised distances (aka Hadamard transforms and therefore the machinery of likelihood models), can be surprisingly high with long sequences (e.g. 4 substitutions per site), before it peaks and relative error rates cause tree recoverability to decline. However, when the model is violated, error rates relative to distances climb much more rapidly, so that recoverability might be hampered at 1 substitution per site or less.

Switching to a codon position partitioned mixture model with independent site models but linked rooted trees/lineage rates, there is another big bump in likelihood (−231,405 vs -237,993 = 6588) relative to the all sites mixture model. The third rate class in particular showed evidence of an inverse relationship of frequencies in rate class 1 versus 2. Root frequency parameter distributions and other parameters agreed across runs, but burn in was slow and one chain failed to converge. Interestingly, the corresponding 4gi model had more trouble achieving high ESS. It seemed one issue there may have been ambiguity of the root and big jumps getting from one root position to another. In terms of the codon-4gi5rf tree, the new tree is much deeper than with the mixture-4gi5rf and there are a few mcc tree changes with flip-flopping strong or very strong support. All but one of these are biologically incorrect. The one plausible group that flip-flops is Euungulata, or Cetartiodactyla + Perissodactyla (pp now >0.9 from ∼0.05) and its local alternative Zooamata = Perissodactyla + Carnivora (pp now ∼0.05 from >0.9). Such a flipflop is also consistent with what was seen in Waddell et al. (1999a,b,c) with different methods of analysis. The unrooted tree remains very much like that in Waddell et al. (1999b,c) which is based on amino acid sequences analysed with ML methods.

This data is large enough to sensibly compare the gtr4gi5rf model to other extensions of the partitioned 4gi model which might be close competitors. One of these, commonly used and available in BEAUti is to unlink the relative rates of substitutions in the three codon positions. While this increases the likelihood by another ∼150 lnL units it comes by adding another 2 x ((2 × 44) – 3)) = 170 free parameters, rather than 3 × 3 × 5 = 45 of the relatively more frugal gtr4gi5rf model. Here, this causes the posterior to largely overlap that of the 5rf model (Figure 8). A further extension is allowing each codon position to have its own relative rates on each edge of the tree and also its own tree. This increased the lnL by about another ∼80, but the posterior was punished by another ∼60 log units. So, for a concatenation of many genes, the new model might be an interesting alternative to simply allowing each gene to have its own edge lengths. If the 4gi5rf model does especially well in such situations, then for unlinked genes, it would suggest that base composition heterogeneity could be exceeding coalescent effects (differing trees and branch lengths) in terms of the overall adjusted fit of the model.

**Figure 8.**
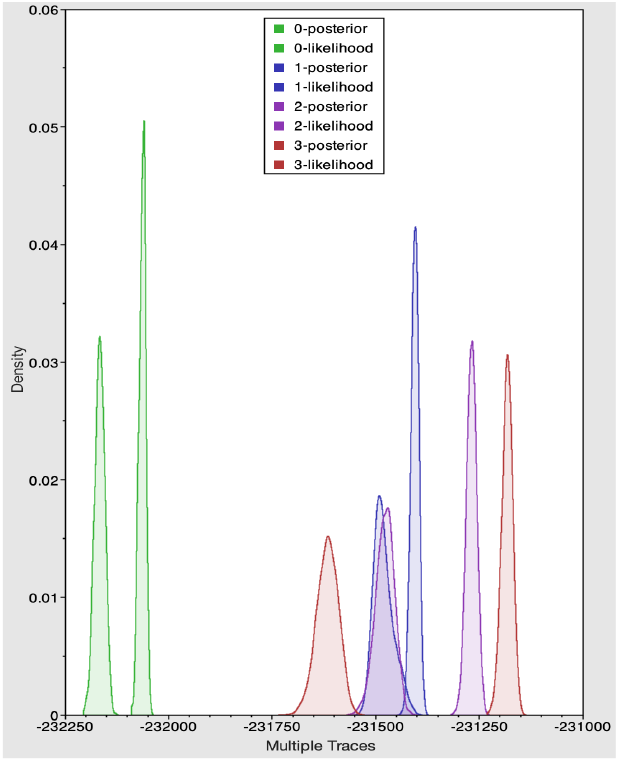
Comparing 4gi5rf and 4gi models with the vertebrate_mtDNA data, with and without codon position partitioning. The models are listed in order of increasing log likelihood and increasing free parameters. Model 0, codon-4gi, Model 1, codon-4gi5rf, Model 2, codon-4gi-unlinked-orc, Model 3, codon-4gi-unlinked-orc+unlinked-trees. The posterior is always to the left and more broad than the corresponding likelihood, and, as with other examples herein, has not been fully normalized.

### Application to short nuclear gene fragments of moderate divergence, ragfib_nuc

In this example the alignment of figure 2 of Waddell and Shelly (2003) (ragfib_nuc) is analysed with bmt4gi5rf and compared to bmt4gi plus the original hky4gi Bayesian analysis (the last using quite different priors, such as, all trees equally likely and branch lengths sampled from an exponential distribution). A look at the NeighborNet visualisation of tests of stationarity and time-reversibility (figure S2.9) suggests that *Tupaia* is strongly divergent in its mutation spectrum, and, yet again, much of the deviation from pairwise symmetry expected by a stationary time reversible model appears to be explained by non-stationarity.

Four chains of bmt4gi_orc_myule using the standard Beauti defaults were run to 10 million steps. Each was checked for convergence visually and the apparent burnin phase ignored. One of the chains malfunctioned, so another was run. Combined post burn in, the ESS of all statistics (posterior ESS 1869) was well over 200 with tree height ESS 741 being the lowest. Eight chains of bt4gi5rf_orc_myule were run to 5 million steps. These all converged quickly to overlapping posterior distributions. The combined posterior ESS was 1882, but tree height and the BICEPS statistic had the lowest ESS (124 and 233, respectively). A plot of the relative posterior probabilities and tree likelihoods are show in figure 9a. Again, there is a substantial improvement in the likelihood for 15 additional free root frequency parameters (about 52lnL units), but in this case it is the simpler model that has the higher posterior probability, probably due to less “evolution” in the data (thus less separation of likelihoods).

**Figure 9.**
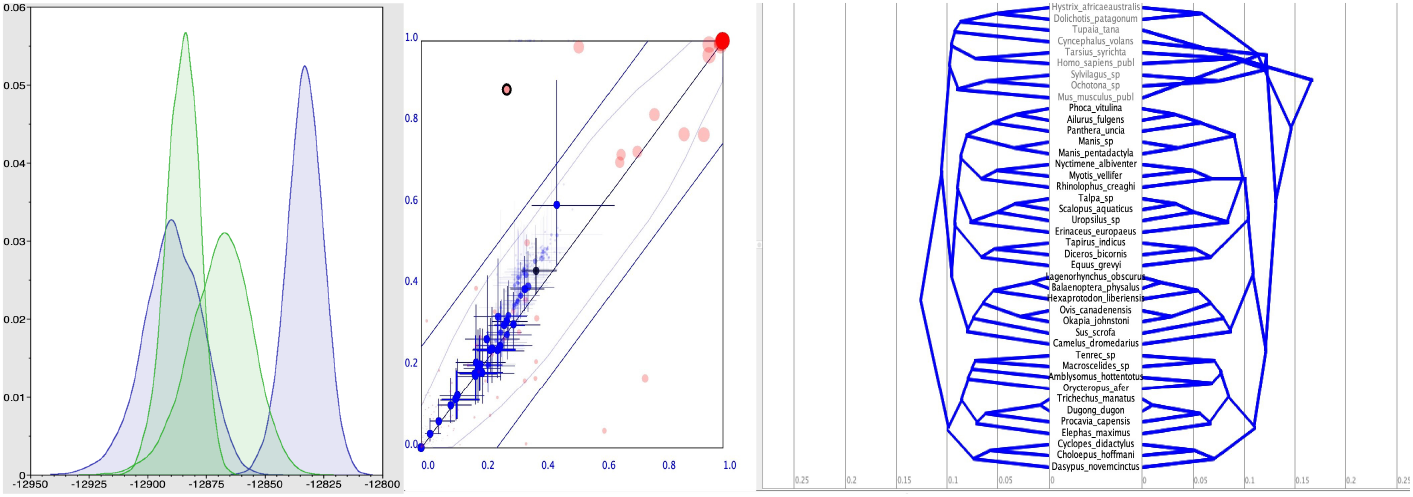
(a) The likelihoods (taller) and posterior probabilities (broader) of the bmt4gi (green) and bmt4gi5rf (blue) models applied to ragfib_nuc. (b) A pairwise plot of clade support values (pink) and divergence time estimates (blue crosses). The higher likelihood but lower posterior model (bmt4gi5rf) appears on the *x*-axis, and the higher posterior model appears on the *y*-axis. (c) A mirrored mcc tree plot with bmt4gi5rf on the left and bmt4gi5rf on the right. The normal arrangement of the simpler model on the left is reversed, because if DensiTree takes the simpler model trees first, and the mirroring then leads to a “bird’s nest” effect.

The mcc trees of these two models are shown in figure 9b. Clearly the 4gi model is having trouble with *Tupaia*, while the new model places the root in a reasonable position, concordant with the clade Atlantogenata (Waddell et al. 1999c, spanning *Tenrec* to *Dasypus* in figure 9c), sister to all other placental mammals. Apart from the root, support for clades showed only a moderate amount of change, with relationships between orders within Laurasiatheria (spanning *Phoca* to *Camelus*) showing generally mediocre support. In terms of divergence times, the 4gi5rf model again produces seemingly more realistic divergence times deeper in the tree, given the complete lack of Mesozoic placental fossils. Highlighted in figure9c is a clade that is almost certainly incorrect with the 4gi model, yet it receives considerable support (pp ∼0.9). The other clade that receives relatively much stronger support in the 4gi model (pp ∼ 0.98) is Atlantogenata. It appropriately receives much more moderate support with the new model (pp ∼ 0.5), given that, under these types of model, there should be relatively little evidence for the exact location of the root. The new model did not greatly differ in its clade support, except near the root, from the unrooted Bayesian analysis in Waddell and Shelly (2003) figure 2.

In terms of the mechanics of the 5rf model and what it is doing, much of this is accommodating just one taxon, *Tupaia*. In this case, the root moves across many nodes, but they are all weakly supported except for the one highlighted in figure 9c. The mechanics remain similar to that seen previously; the root frequencies that matter most in terms of the tree (those of the fastest evolving sites), start somewhat beyond any given taxon and the stationary frequencies place in an opposite direction beyond that of the most derived and deviant taxa (*Tupaia*). Here, however, there was a marked contrast in the root frequencies of the second fastest rate class compared to the fastest. These sites are is markedly more important given a lesser degree of rate inequality. This suggests that between the evolving sites there are other patterns of heterogeneity that the free root frequencies are responding to.

## Discussion

Applied to simulated data of a realistic length generated from a set of broad priors, the model is shown to be able to recover all parameters with close to the accuracy predicted by the model (a 95% credibility interval). That is, the 4gi5rf model implementation has achieved practical identifiability of parameters, including the tree itself. Both these simulations and theoretical understanding do caution a few instances where practical identifiability can break down. One of these is distinguishing the invariant sites from the slowest evolving sites (e.g., the slowest quartile when using a four category gamma approximation, but also occurring with a true gamma distribution). By allowing each rate category of sites to have its own root base composition (which is then, effectively, its only base composition if at a very slow or zero rate), this too invites diminution of practical identifiability. This emerges in the form of broad distributions on the posteriors of the exact frequency of each base, and, perhaps the posteriors of these two categories mirroring each other (e.g. if one has a high frequency of A, the other takes on low A). With these caveats in mind, however, the model and its implementation appear a useful extension of 4gi models.

Applied to a the widely used primate_mtDNA data, the model shows up well in comparison to major steps in the development of likelihood models over time, from Jukes Cantor to general time reversible with invariant sites and gamma distributed rates, that is gtr4gi. It does this in terms of not only its ability to strongly increase the likelihood and posterior probabilities, but it also alters support values and divergence times of clades more so than many of these other important steps in evolutionary model development. It might even be tackling a problem that can be particularly acute for the gtr4gi model when base compositions are non-stationary, near the root in particular. In such situations “long edges repel” (Waddell 1995) can be a serious problem. The model also performed well on this test data with multiple time calibrations and monophyletic constraints in place. These “force” reconstructions to avoid embarrassments like “Jurassic Monkey,” but in so doing make many methods behave much more similarly. It was necessary to remove constraints to understand what substitution models are doing with real data.

The gtr4gi5rf model (and its sub-models) can counter overestimation of path lengths and hence rates due to non-stationary base composition. It does this most effectively near the root, but the “gradient” of non-stationarity it sets up from root to the most derived tip can have effects further down the tree. This seemed to have a mostly positive effect on the example data, where re-rooting was reasonable, and often corrected errors near the root introduced with the use of the gtr4gi model. Note, over estimation of certain paths due to non-stationary evolution can also lead to, for example, the anti-Felsenstein tree of long branches together repelling each other so a Felsenstein tree is incorrectly inferred by stationary models (Waddell 1995). Another way of discussing this type of effect is as a combination of rate/time plus process impacting total relative branch lengths (Lockhart and Steel 2005). In some of the example data analyses, the 4gi models seem to overestimating the longest and/or deepest branch lengths due to non-stationarity, with corresponding effects on the inferred location of the root.

A sobering result of these investigations with real data is that neither relative likelihood nor posterior probabilities are a reliable guide to the improvement of key phylogenetic parameters. Indeed it was seen that along a series of major steps in the improvements of the likelihood, the key parameters, clades, along with their estimated probability of being correct, and relative node heights, have often gotten better, worse or not changed. The popular gtr4gi produced some of the most biologically troubling results on the real data. Also, on real data sets we see multiple examples where posterior proportion support values of clades really flipflop with changes to the analysis; always a concern. Accordingly, many of these models do not seem robust in their finer details, including giving extreme posterior support values when the data fail to fit the models assumptions.

Similar concerns apply to relative divergence time estimates, where we saw quite different estimated node heights deep in the tree, including when using nuclear sequences. This has direct implications for unsettled questions, such as how old are the orders and superorders of placental mammals are (Waddell 1999c, Waddell et al. 2001, Kitazoe et al. 2007) and in the case of superordinal clades within Laurasiatheria, which ones are actually correct. Indeed how can we have any particular preference when the likelihood and posterior probability of current models seem a poor guide to correctly estimating fundamental parameters when they differ?

A continuing issue is if or how to partition nucleotide data and/or whether to use a mixture model, or some combination. Thus the comparable likelihood of the 5rf model (via 15 extra parameters) as an alternative to allowing each branch of a tree to have its own length per rate class (4n-4 extra parameters) warrants further evaluation of this model for concatenated alignments, with the former being a potentially much more parsimonious solution.

In this article we explored the utility of allowing just the root to adopt its own base frequency vector for each rate class. This seems to help near the root, but the effect down the tree is rapidly diluted. It seems that a five rate class model is appropriate at the root, but base frequency vectors defining the transition matrices need to change when the transition matrices change within the tree, something implied in the supplementary analyses of the all biological data considered.

Tackling base composition shifts has long been identified as a major issue. Now it is perhaps the major issue facing neutral nucleotide models of molecular evolution applied in the majority of situations beyond the last few thousand generations. There remain well-known data sets where all current models fail. One of these is a vertebrate_mtDNA alignment. The gtr4gi5rf model seemed to hardly make a dent on the problems with that tree. That might well be indicative of how difficult it is to confidently resolve really deep divergences in the tree of life, such amongst early eukaryotes or bacterial lineages. It corroborates the analyses in Waddell et al. (1999b, 2001) that some of the mitochondrial amino-acid coding sequences are so difficult to model, that they should to be excluded to obtain final robust results.

The current model can be thought of also as exploring the root contribution of a more complex non-reversible and non-stationary model, that also adjusts at the same time for base composition heterogeneity across rate classes. Further down the tree, it seems important to implement a model that frugally allows the rate matrix to change at certain points (Foster 2004). The current model does particularly well at modelling the slowest rate categories. Although 15 extra parameters is frugal in comparison to partitioning schemes that frequently add in 2n-2 extra branch rates (per partition), it could also be interesting in future to explore merging certain rate class parameters.

## Software availability

An open source implementation is available under the LGPL 3.0 license in the beastbooster package for BEAST 2, available from https://github.com/rbouckaert/beastbooster. BEAST2 is open source and freely available from http://beast2.org/.

## Data availability

data for the well calibrated simulation studies as well as the BEAST XML files for the empirical data studies are available from https://github.com/rbouckaert/baseCompositionManuscript

## Acknowledgements

This work was partly supported by NIH grant 5R01LM008626 to PJW.

## Notes

### Competing Interest Statement

The authors have declared no competing interest.

