## Supplement0 for "An independent base composition of each rate class for improved likelihood-based phylogeny estimation; the 5rf model"

Supplement 0. Practical Identifiability via simulations

Presented below are the results of 100 DNA alignments of sequence length 500, with generating parameters sampled from the priors listed below, then analysed with Beast2.5 using the hky4gi5rf model.

The priors used to both simulate and to analyse the data were:
Yule tree prior with birth rate distribution uniform, lower bound = 5 and upper bound = 6.
Root age prior a uniform distribution, lower = 0.25 and upper = 1.0.
All frequency prior vectors distributed as Dirichlet, alpha = 4.0, 4.0, 4.0, 4.0.
PropInvariable has prior beta distributed with alpha = 1.0 and beta = 4.0.
Gamma shape prior is exponential with mean = 1.0.
The hky kappa value has a log normal prior with mean, M = 1.0, and standard deviation, S = 1.25.

(see next page)

### Coverage calculations

prior sample: ../truth.log
posterior samples: summary with 100 runs so coverage should be from 91 to 99

#### TreeHeight

Coverage: 91 Mean: 43 ESS (mean/min): 777/479

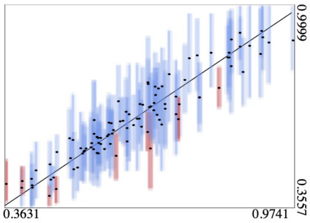


#### YuleModel

Coverage: 96 Mean: 52 ESS (mean/min): 700/257

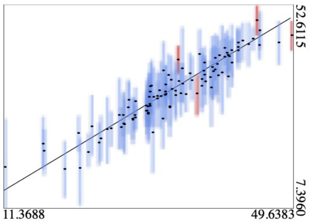


#### birthRate

Coverage: 93 Mean: 49 ESS (mean/min): 840/566

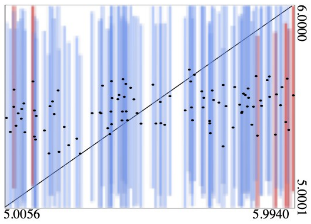


#### kappa

Coverage: 94 Mean: 49 ESS (mean/min): 718/484

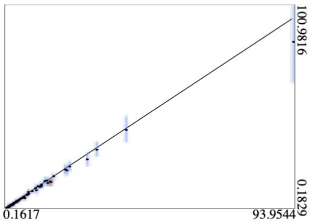


#### proportionInvariant

Coverage: 91 Mean: 47 ESS (mean/min): 436/35

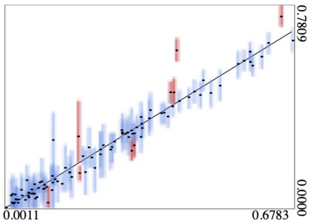


#### gammaShape

Coverage: 91 Mean: 50 ESS (mean/min): 532/22

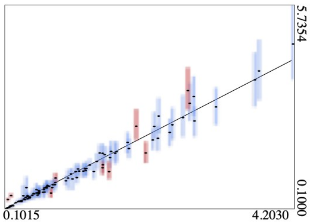


#### freqParameter.1

Coverage: 94 Mean: 50 ESS (mean/min): 839/536

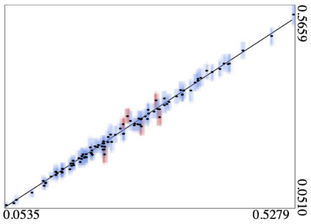


#### freqParameter.2

Coverage: 94 Mean: 41 ESS (mean/min): 836/648

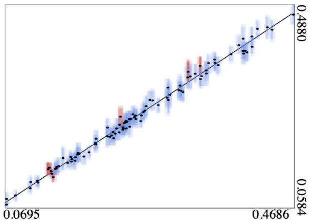


#### freqParameter.3

Coverage: 93 Mean: 54 ESS (mean/min): 833/513

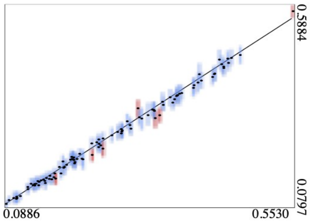


#### freqParameter.4

Coverage: 94 Mean: 52 ESS (mean/min): 826/473

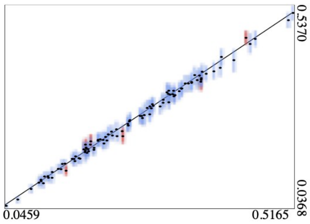


#### freqParameter5.1

Coverage: 92 Mean: 58 ESS (mean/min): 791/290

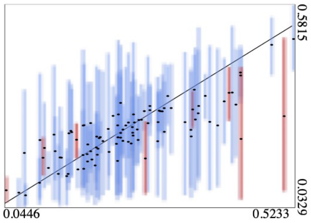


#### freqParameter5.2

Coverage: 88 Mean: 47 ESS (mean/min): 778/294

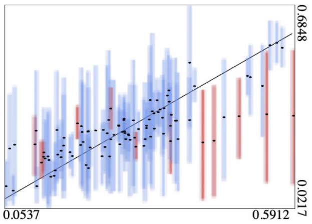


#### freqParameter5.3

Coverage: 92 Mean: 42 ESS (mean/min): 786/268

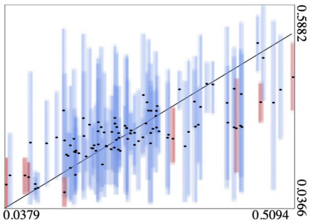


#### freqParameter5.4

Coverage: 92 Mean: 38 ESS (mean/min): 780/184

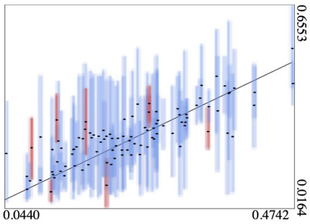


#### freqParameter1.1

Coverage: 94 Mean: 42 ESS (mean/min): 767/71

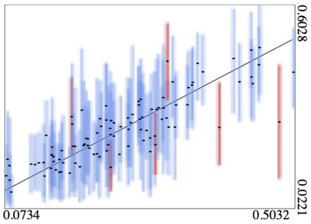


#### freqParameter1.2

Coverage: 92 Mean: 53 ESS (mean/min): 723/118

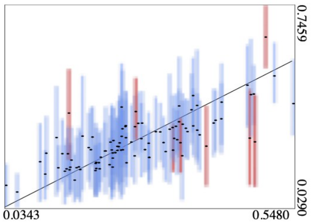


#### freqParameter1.3

Coverage: 93 Mean: 47 ESS (mean/min): 759/147

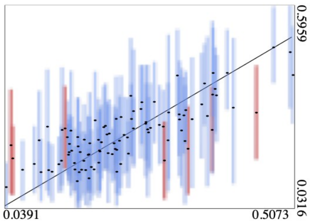


#### freqParameter1.4

Coverage: 93 Mean: 55 ESS (mean/min): 740/98

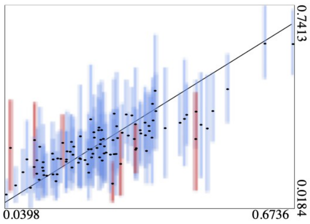


#### freqParameter2.1

Coverage: 93 Mean: 48 ESS (mean/min): 763/49

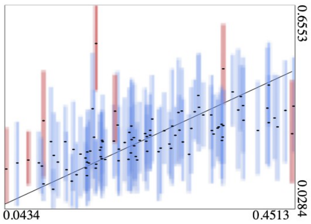


#### freqParameter2.2

Coverage: 93 Mean: 46 ESS (mean/min): 798/184

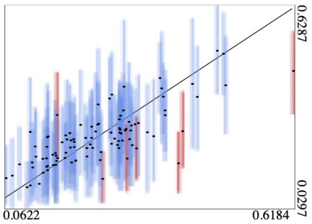


#### freqParameter2.3

Coverage: 95 Mean: 59 ESS (mean/min): 778/228

#### freqParameter2.4

Coverage: 90 Mean: 48 ESS (mean/min): 776/77

#### freqParameter3.1

Coverage: 96 Mean: 54 ESS (mean/min): 826/217

#### freqParameter3.2

Coverage: 94 Mean: 40 ESS (mean/min): 824/610

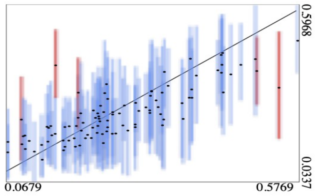


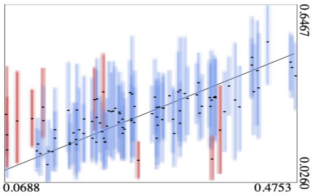


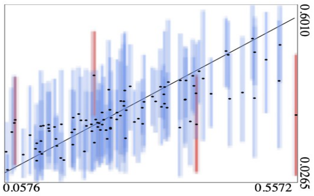


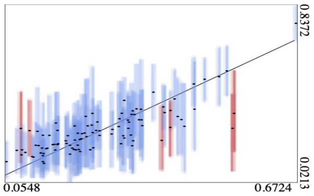


**freqParameter3.3**

Coverage: 98 Mean: 48 ESS (mean/min): 805/273

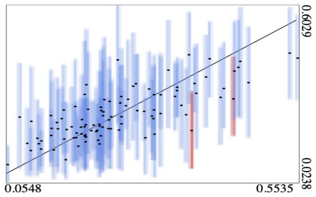


**freqParameter3.4**

Coverage: 95 Mean: 46 ESS (mean/min): 826/435

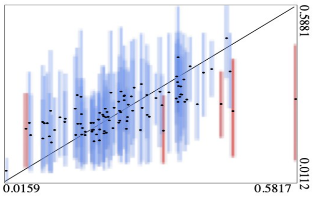


**freqParameter4.1**

Coverage: 92 Mean: 48 ESS (mean/min): 842/448

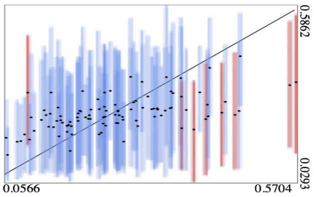


**freqParameter4.2**

Coverage: 95 Mean: 49 ESS (mean/min): 834/460

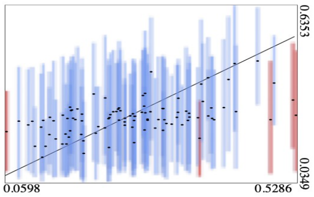


**freqParameter4.3**

Coverage: 92 Mean: 47 ESS (mean/min): 841/582

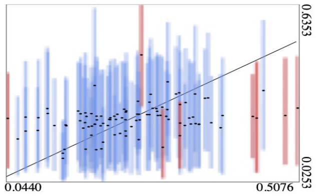


**freqParameter4.4**

Coverage: 95 Mean: 45 ESS (mean/min): 838/558

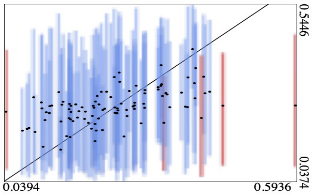


* marked ESSs indicate one or more ESS estimates are invalid. Unmarked ESSs indicate all estimates are valid.
