## Supplement2 for "An independent base composition of each rate class for improved likelihood-based phylogeny estimation; the 5rf model"

Supplement 2: Analyzing base composition and transition matrix shifts using INTEROGATE

This supplement makes a series of tests (and visualisations of test results) for base composition stationarity and deviation from a time-reversible evolutionary process. Results are shown in turn for the primate_mtDNA, vertebrate_mtDNA and then ragfib_nuc datasets.

The first results show use the generalized sum of squares pairwise test (SS5) of the stationarity of base composition (Tavere 1986). Under the null hypothesis of stationarity, the test statistic asymptotically follows a chi-square distribution (given sufficient data) with 3 degrees of freedom. Accordingly, a value of 7.82 or greater has a probability of less than 0.05 under the null hypothesis of stationary base composition. In order to ensure closer convergence to asymptotic conditions, and to avoid non-invertible estimated covariance matrices, a grouping of base frequency differences is performed, so that all expected values are 5 or greater (using the FreqNuc program of INTEROGATE, Waddell et al. 2004). Many of the individual pairs of taxa are significantly non-stationary.

The symmetry of the pairwise divergence matrix between all pairs of taxa was also assessed. The metric that is used is twice the log likelihood of the ratio of the likelihood of the pairwise site pattern frequency matrix (divergence) matrix when it is unconstrained versus when it is symmetricized; the G2Sym test. This test has six degrees of freedom, and, with sufficient expected substitutions between a pair of taxa, closely follows a chi-square distribution. Accordingly, a value >12.59 has a probability of less than 0.05 under the null hypothesis that the pairwise divergence matrix is symmetric; when symmetry is broken the implied evolutionary process is not time reversible.

A complete set of pairwise test results for a common multiple sequence alignments were visualized with NeighborNet (Bryant and Moulton 2004) as implemented in SplitsTree4 (Huson and Bryant 2006). In this case, the pairwise test statistic is used as the distance measure. These vizualizations were compared to NJ trees also using SplitsTree4. In all these cases the NeighborNet had a noticeably better fit, consistent with the appearance of "boxiness" in its graph. The results for the SS5 and G2Sym tests, respectively, on primate_mtDNA are shown in figure S2.1 and S2.2. As can be seen, many pairs of taxa are significantly non-stationary with respect to each other and also non-symmetric in their divergence. Clearly, different groups of primates and outgroup taxa violate the assumptions of stationarity and reversibility in quite different ways and to different degrees.

Note also that the taxa *Lepilemur* and *Daubentonia*, two labile taxa in the tree analyses of the main text, are surrounded by taxa moving in quite different directions regarding shifting substitution spectra. The shifts in the base composition of taxa involved in the near tricotomy of Perissodactyla, Cetartiodactyla and Carnivora, are much less pronounced, yet the time durations involved are longer (e.g., ~15-55 mya vs ~55-100 mya).

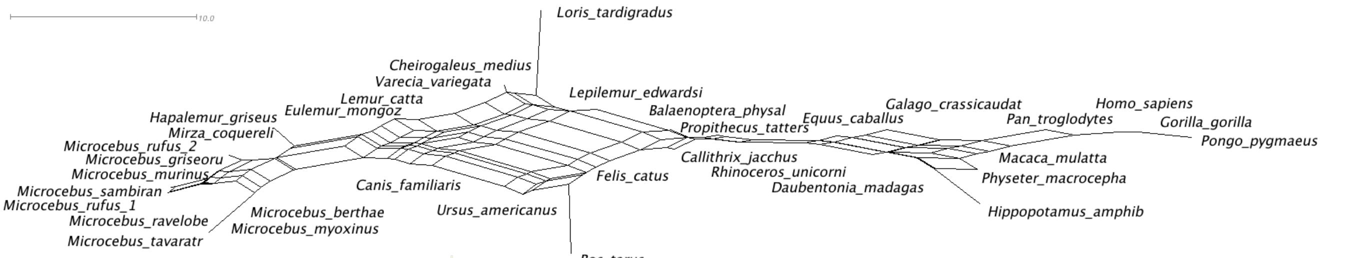


Figure S2.1. A Neighbor Net visualization of all GLS SS5 pairwise tests of base-composition stationarity applied to the primate_mtDNA multiple alignment. The percentages of A, C, G and T for *Microcebus rufus* 1 are 31 25 14 31, while those of *Pongo* are 29 34 13 25. Additionally, those of *Loris* are 33 27 12 27, and those of *Bos* are 32 28 14 26. Thus, the major left to right trend in the network is due mostly to increasing C and decreasing T, while the trend from *Loris* to *Bos* correlated with G increasing. When fitting the gtr4gi5rf model to these sequences, the strongly favored stationary composition was 20 53 3 24. This is well outside the observed composition of any of these taxa, but it is in the direction the anthropoids are headed relative to *Microcebus*. In contrast the gtr4gi model gave stationary frequencies of 36 37 6 21, which are much closer to the observed overall frequencies (although these include invariant sites, as this model does not distinguish their frequencies).

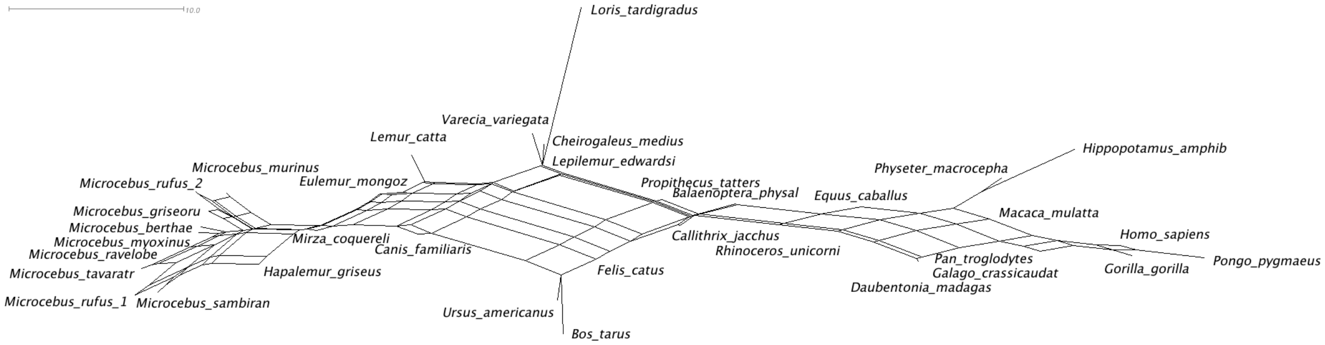


Figure S2.2. A NeighborNet visualization of the pairwise distances induced by the G2Sym test applied to the primate_mtDNA data.

Figures S2.3 and 4 show the comparative test results after translation to amino acid sequences. These follow a similar pattern to the raw nucleotide sequences, but with a lot more idiosyncrasy of individual taxa (shown by terminal edges generally being much longer). This suggests that base composition shifts are a major factor in amino acid changes that are non-reversible and non-stationary. Note also that to be significant, the test statistics need to be markedly larger (e.g. 19 versus 3 degrees of freedom), so some of the idiosyncrasy (e.g., expressing as relatively longer terminal branches) in the graph is expected due to random fluctuations.

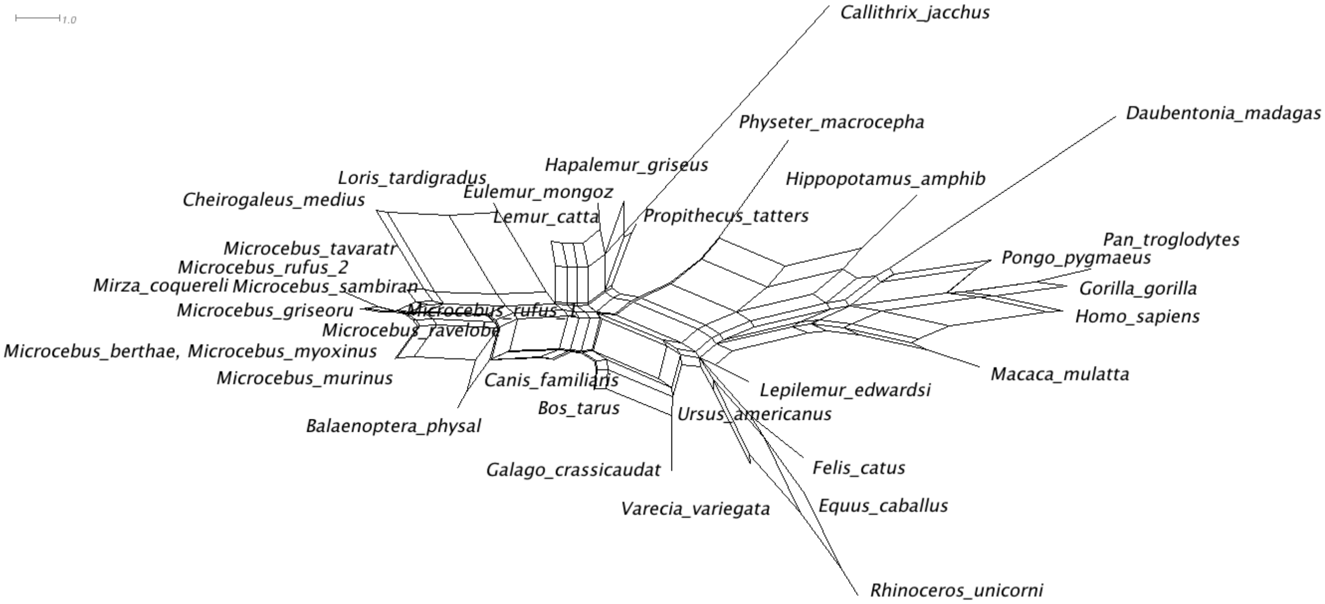


Figure S2.3. A NeighborNet visualization of the pairwise distances induced by a SS5 pairwise tests of amino acid composition stationarity for primate_mtDNA.

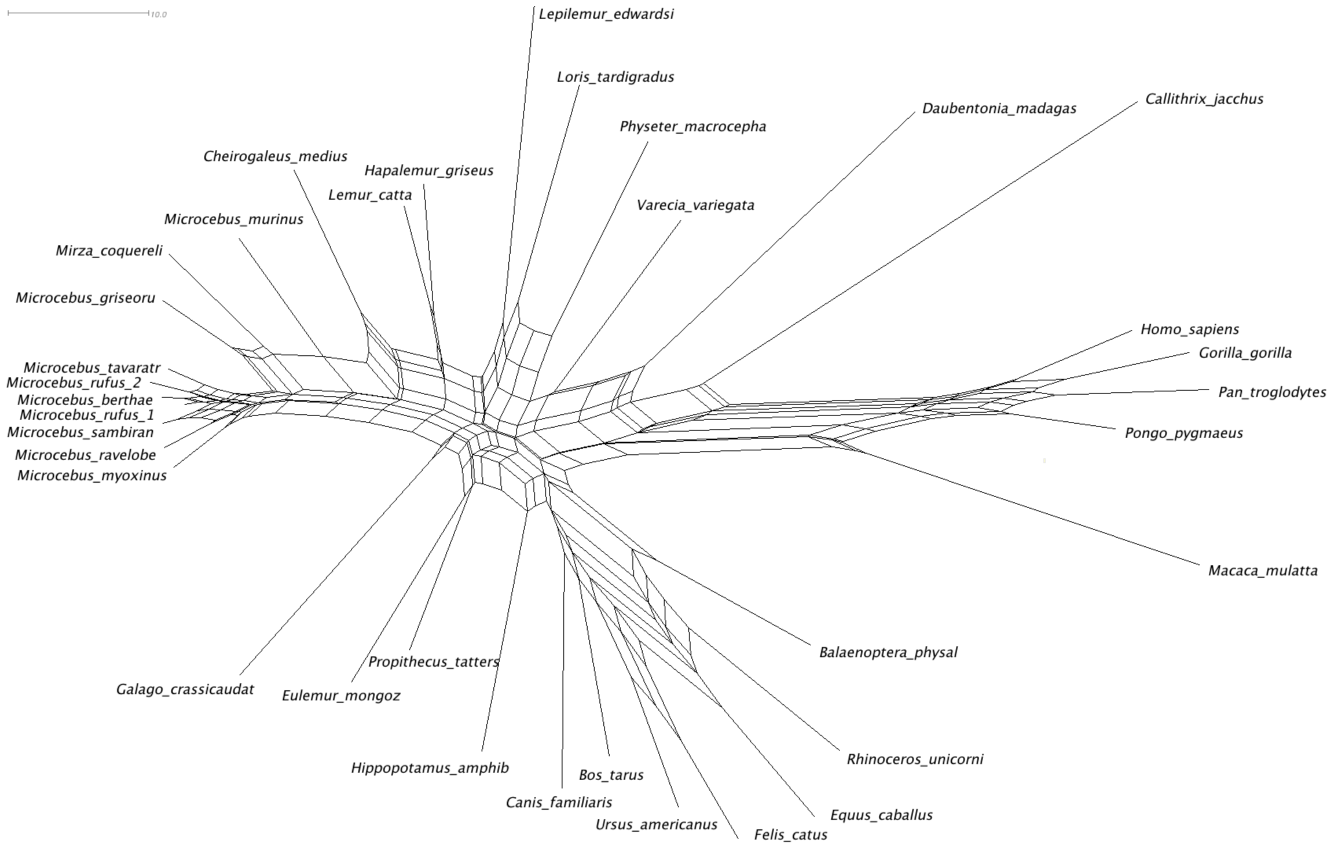


Figure S2.4. A NeighborNet visualization of the pairwise distances induced by the G2Sym tests applied to the amino acid translation of primate_mtDNA.

It is also useful to also look at how these tests apply as first, second and third coding positions are considered in turn (S2.5-7). In terms of total branch lengths measured by parsimony on a tree (the NJ tree of all sites with Hamming distances), the first position sites have total branch lengths of 1255, the second position sites 416, and the third position sites 4683. The model based differences in substation rate are even larger, for example, the gtr4gi5f model suggests relative rates of 0.32, 0.07 and 2.62. However, since the stationarity tests here are measured on the observed data, as are parsimony changes, these may be the better guide to power for tests in each data partition.

One important thing to note in the following six visualizations is that the test results are most consistent with deviations from stationarity being most tree-like for third position sites, markedly less so for first position sites and least similar for second position sites. The SS5 results for second positions strongly emphasize that the anthropoids and *Daubentonia* are operating with a substitution bias affecting amino-acid changes, that is unlike any other mammals in this data. A similar trend is seen with regards to the similarity of the stationarity and symmetry tests. Indeed, when results are contrasted between third position sites (with the overall weakest functional selection) versus the second position sites (with the strongest functional selection), it seems that symmetry in the slower changing second position sites is being broken due to factors unlike the more recent base composition shifts best observed in the third position sites. However, there is a considerable loss of power in the tests due to much less evolution at second position sites so, the terminal lengths in particular, are expected to increase due to idiosyncratic random stochastic fluctuations.

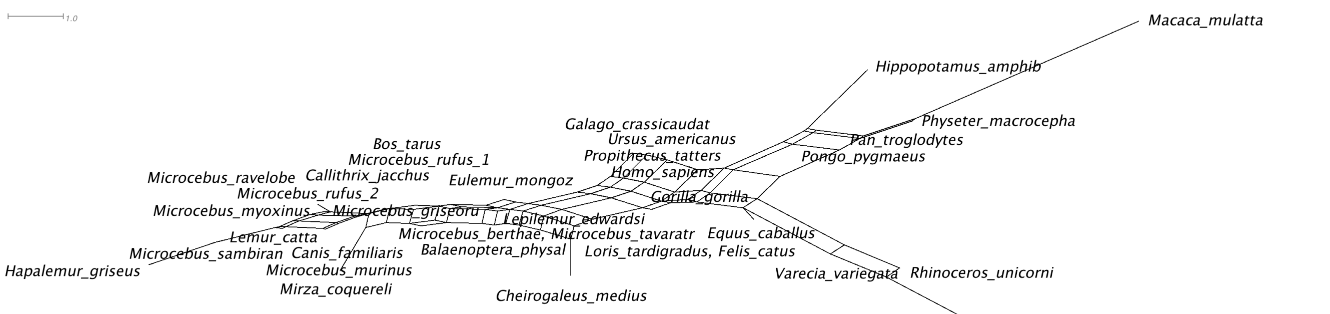


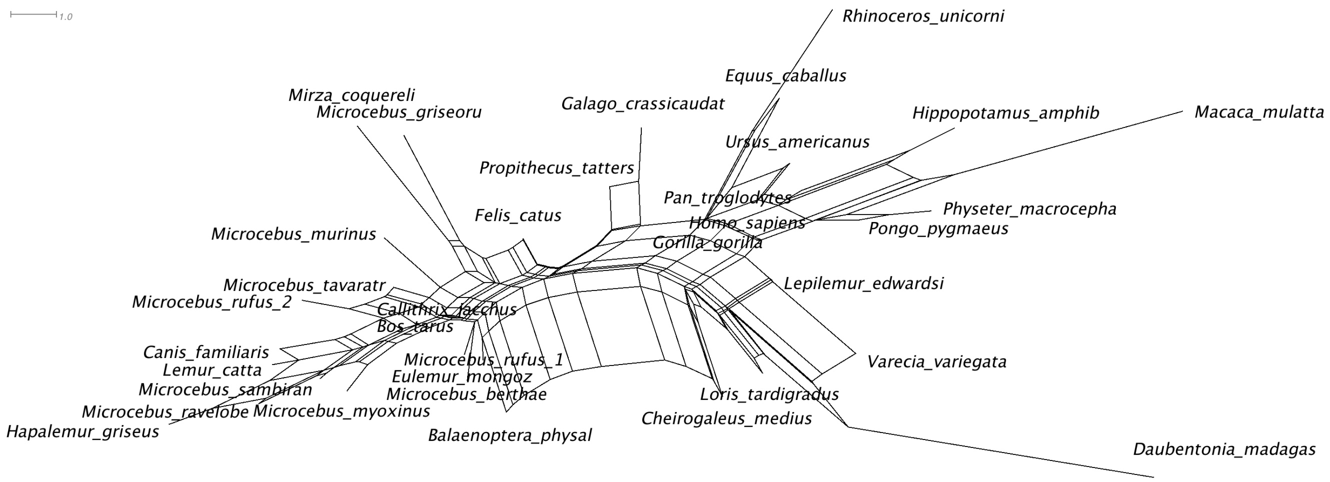


Figure S2.5. The SS5 then G2Sym NeighborNet visualized test results for the mtDNA_primates first position sites.

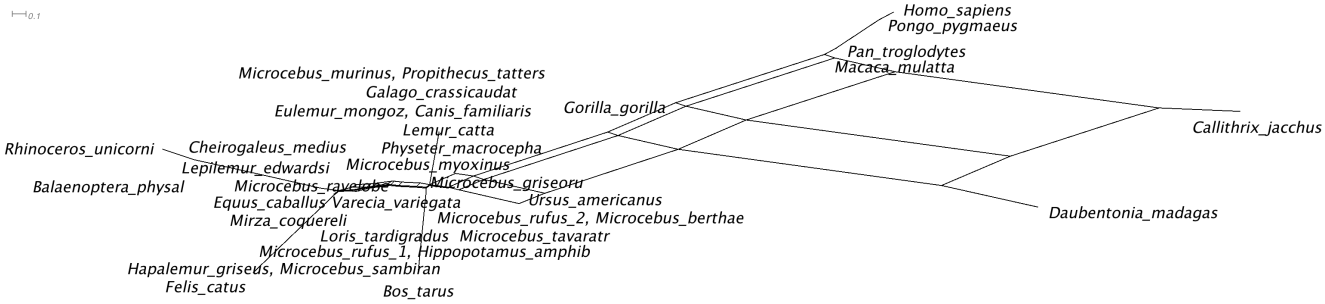

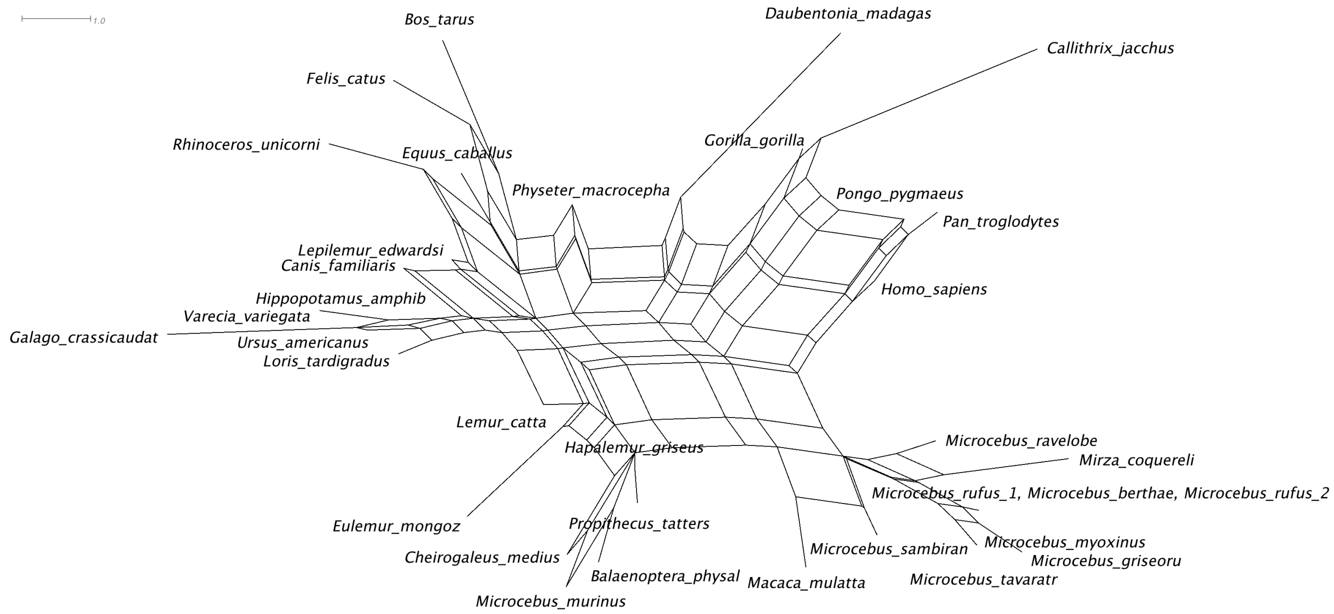


Figure S2.6. The SS5 then G2Sym NeighborNet visualized test results for the mtDNA_primates second position sites.

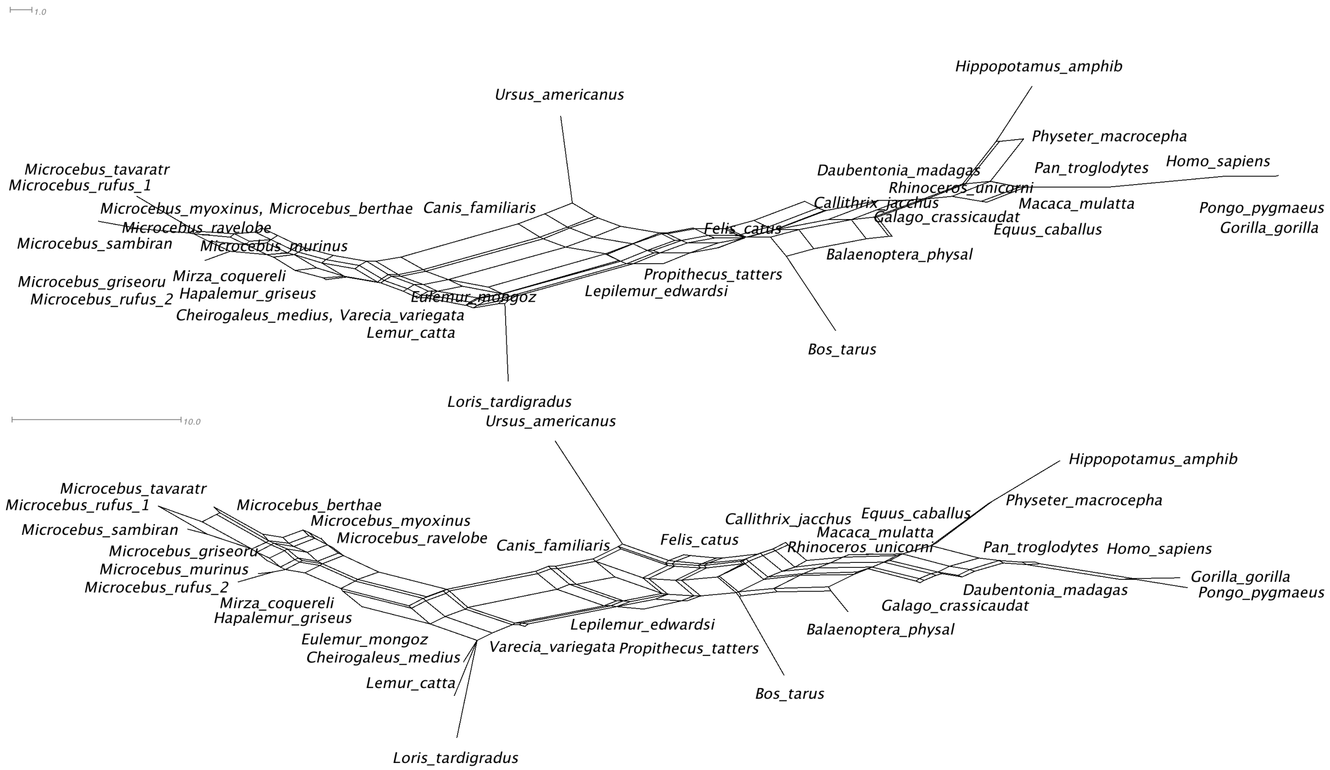


Figure S2.7. The SS5 then G2Sym NeighborNet visualized test results for the primate_mtDNA third position sites.

For the 12 protein coding vertebrate_mtDNA data the extremes of non-stationarity and non-symmetry are larger, but the differences within Primates are still quite extreme even in this context (figure S2.8). Here the extremes are hedgehog on the left with A, C, G, and T of 31 21 12 36, while the duck on the right is 28 35 15 23. So, again, the main left to right trend is dominated by increasing C and decreasing T. The trout has composition 26 30 16 28 while the Coelocanth has composition 33 28 14 25. This suggests a trend of decreasing A from top to bottom. Again the symmetry tests results are very similar to those of base composition, suggesting that drifting base composition explains most of the loss of symmetry. The 4gi5rf model applied to this data favors a stationary frequency of 23 46 3 28. This suggests a trend combining some aspects of the trout and the duck. Very broadly, this is like the stationary frequencies for the primate_mtDNA data, but, in this case, it does not result in a major rerooting with the new 4gi5rf model. The standard 4gi model has stationary frequencies of 38 33 6 23.

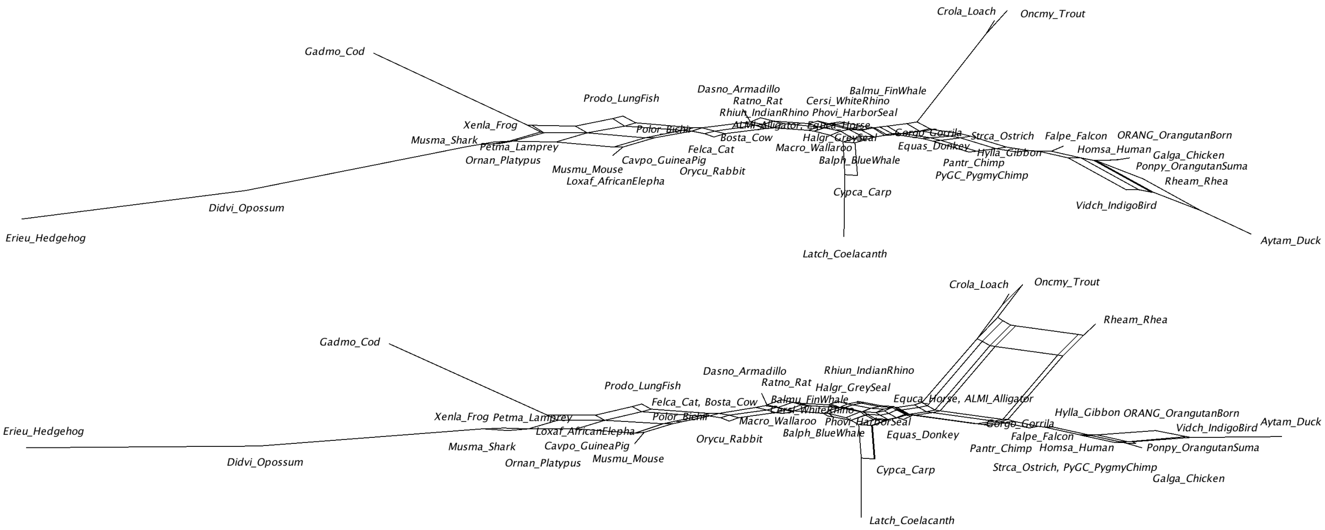


Figure S2.8_1. The SS5 then G2Sym NeighborNet visualized test results for the vertebrate_mtDNA sites.

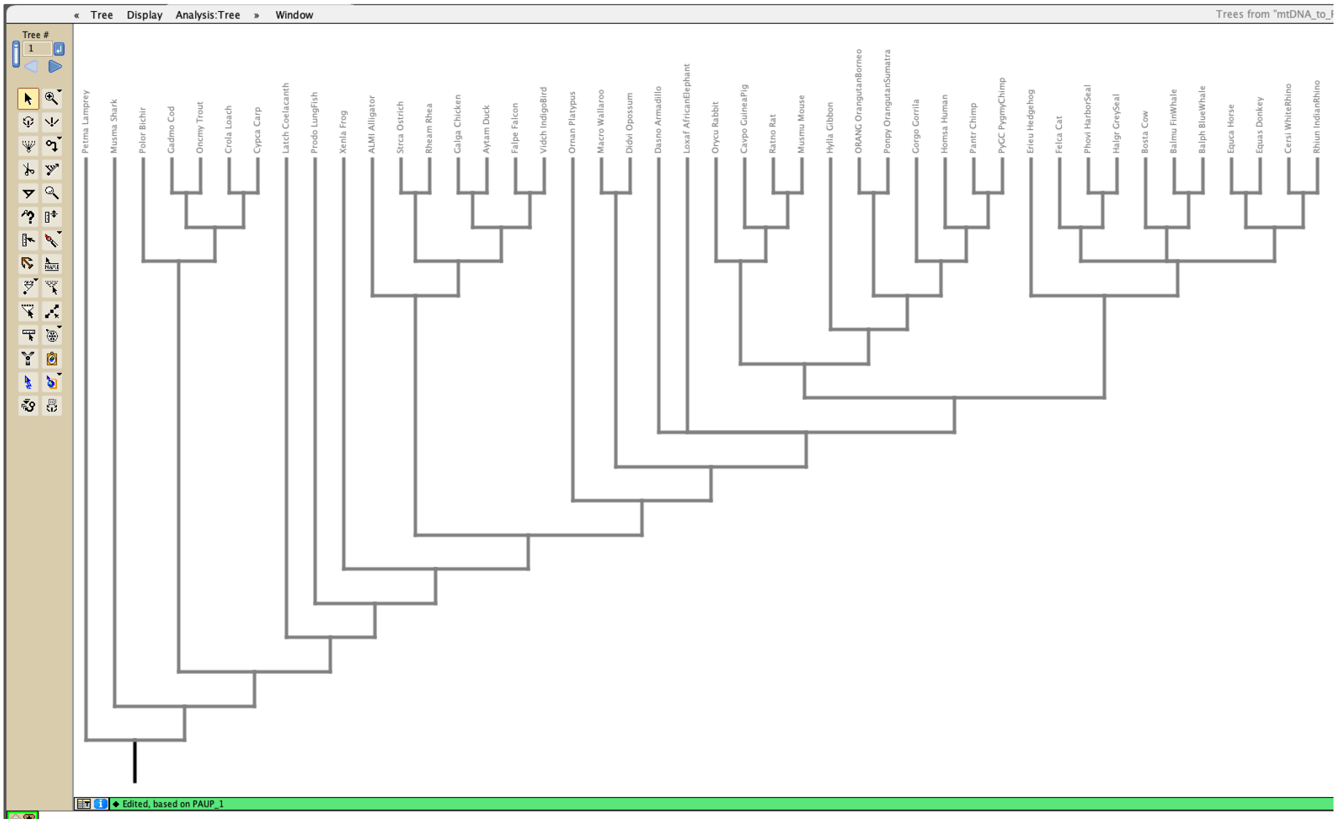

Figure S2.8_2 The expected true tree for the vertebrate_mtDNA. The unresolved relationship reflects the balance of current evidence, with as yet, no evidence convincingly resolving this part of the tree.

### Assessing base composition shifts in the ragfib alignment

Figure S2.9 shows the results of our tests for the nuclear ragfib_nuc alignment. The standout feature here is the strong base composition shift in *Tupaia*. In addition, shifts in the G2Sym test results strongly mirror those of the SS5 results, suggesting that most of the non-symmetry in pairwise divergence (movement away from a time reversible model) is due to the substitution frequency spectrum changing.

In this figure the composition of *Tupaia* is 26 30 28 16, that of *Orycterpus* 30 24 22 23, that of *Panthera* 29 24 25 22 and that of *Sylvilagus* 26 24 26 24. This suggests a trend in increasing A and T, with decreasing C and G from left to right and a further drop in A from top to bottom. The root frequencies of the highest and second highest rate classes of the 4gi5rf model are 23 9 27 41 and 19 36 12 32, while the stationary frequencies are 15 44 22 19. The stationary frequencies of the 4gi model are 26 29 25 20. It seems the the 4gi5rf model is largely adapting to the *Tupaia* sequence by starting the fastest evolving sites at a frequency opposite to *Tupaia* and tilting the stationary frequencies well beyond the trend towards *Tupaia.* This adjustment of parameters to increase likelihood is expected and is seen in the other examples.

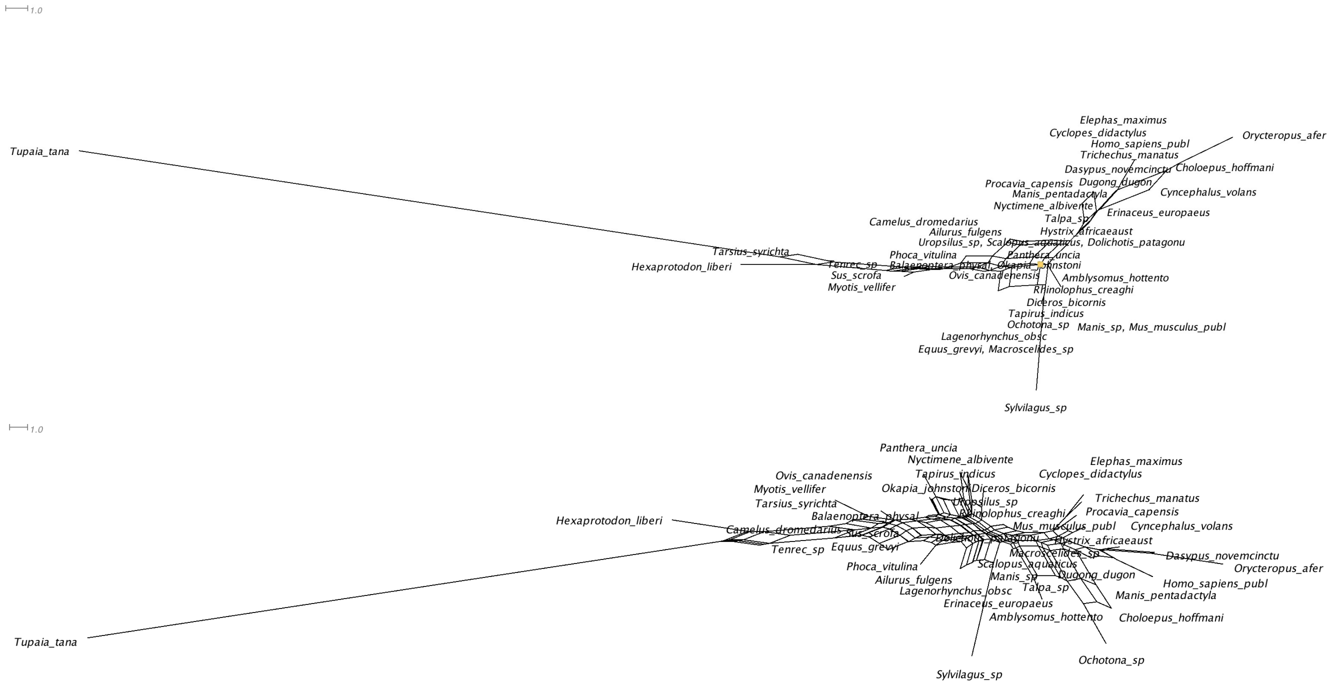


Figure S2.9. The SS5 then G2Sym NeighborNet visualized test results for the ragfib_nuc alignment.
