## Supplement1 for "An independent base composition of each rate class for improved likelihood-based phylogeny estimation; the 5rf model"

Supplement 1: DensiTree2 comparisons for the series of models applied to primate_mtDNA data.

These figures illustrate changes in the trees for the series of seven notable steps in likelihood models. These comparisons are made from the set of trees from the older and, in this case, lower likelihood model, to the next model up in the series unless otherwise noted. The most dramatic changes in clade support are found near the top left and bottom right corners of the *x*-*y* plots on the left, and one of these will usually be highlighted.

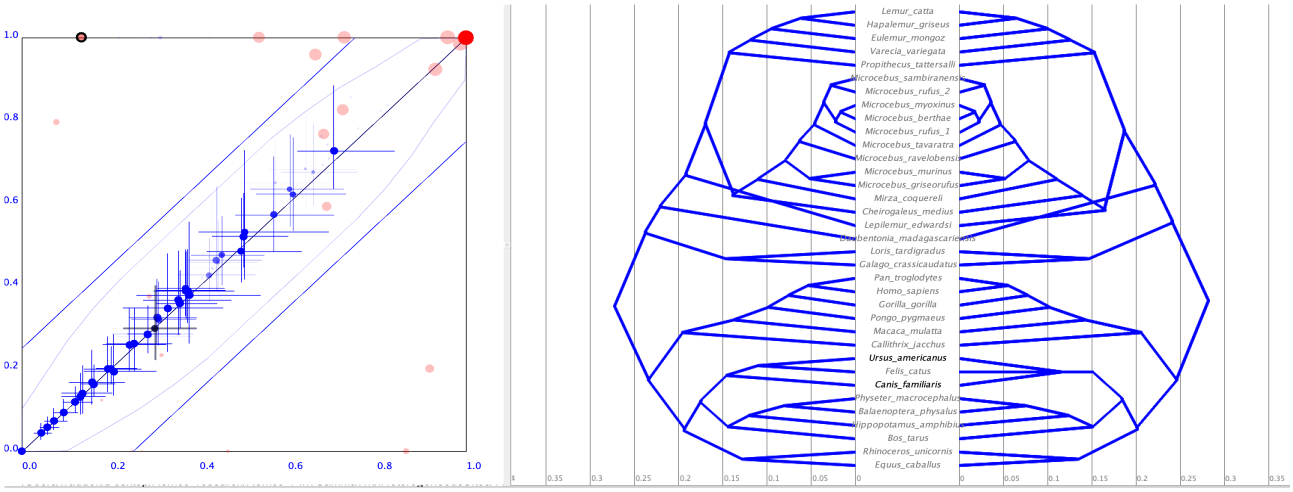


Figure S1.1. (a, leftmost) A pairwise *x*-*y* plot of clade support values (pink) and divergence time estimates (blue crosses). The lower likelihood model (jc) appears on the *x*-axis, and the next higher likelihood model (k2) appears on the *y*-axis. (b) A mirrored mcc tree plot with jc on the left and k2 on the right.

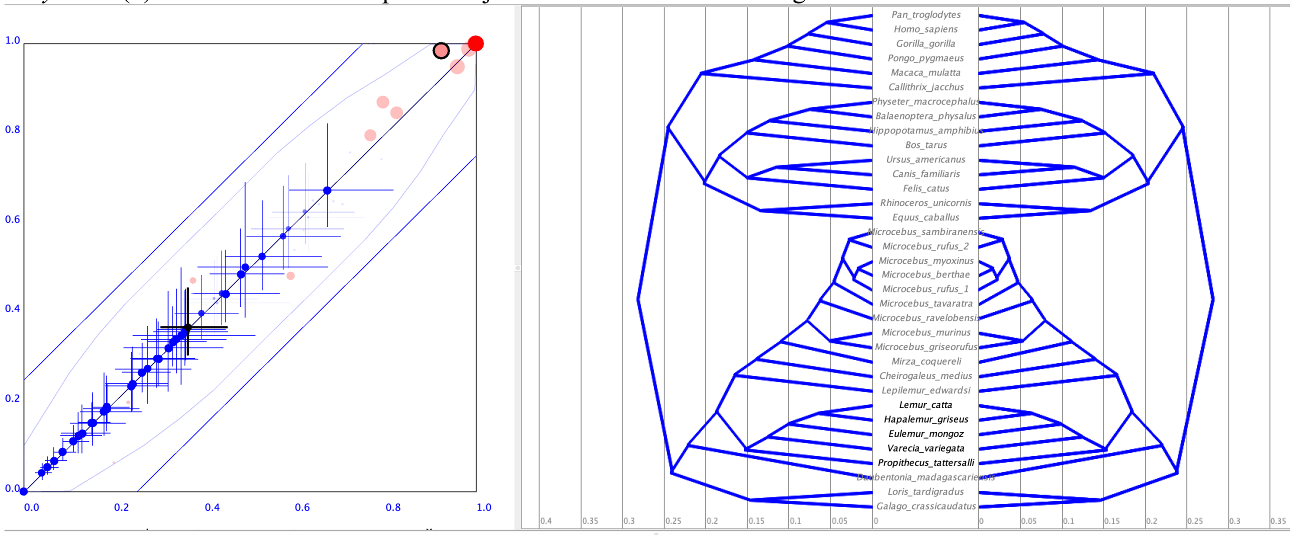


Figure S1.2. (a) A pairwise plot of clade support values (pink) and divergence time estimates (blue crosses). The lower likelihood model (k2) appears on the *x*-axis, and the next higher likelihood model (hky) appears on the *y*-axis. (b) A mirrored mcc tree plot with k2 on the left and hky on the right. Note, despite an appreciable jump in likelihood (main text figure 2), the trees and clade support are very similar.

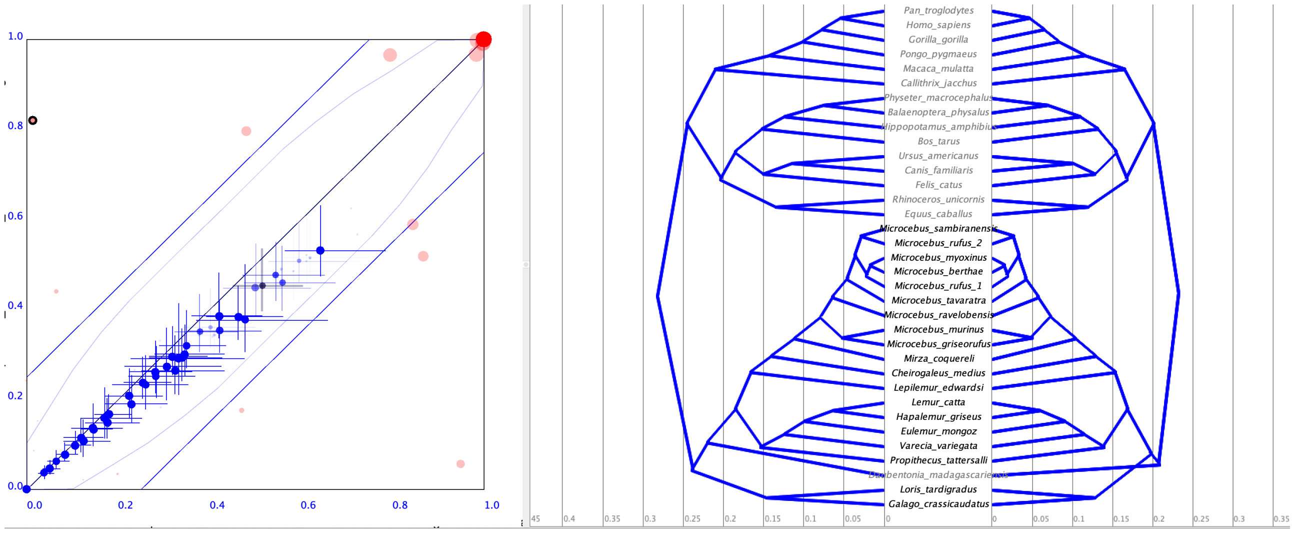


Figure S1.3. (a) A pairwise plot of clade support values (pink) and divergence time estimates (blue crosses). The lower likelihood model (hky) appears on the *x*-axis, and the next higher likelihood model (gtr) appears on the *y*-axis. (b) A mirrored mcc tree plot with hky on the left and gtr on the right.

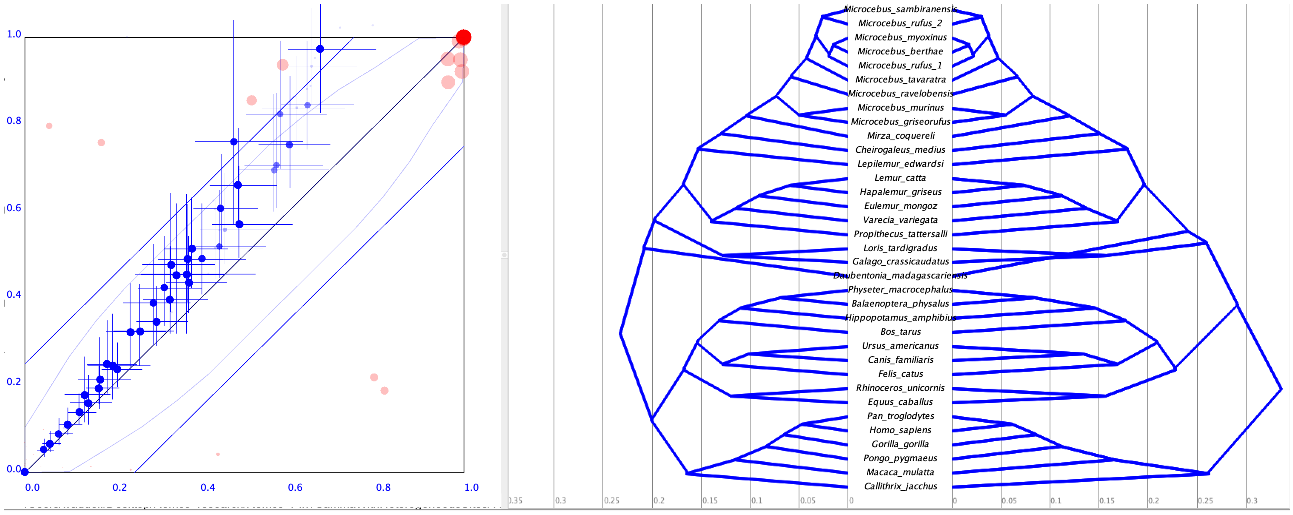


Figure S1.4. (a) A pairwise plot of clade support values (pink) and divergence time estimates (blue crosses). The lower likelihood model (gtr) appears on the *x*-axis, and the next higher likelihood model (gtri) appears on the *y*-axis. (b) A mirrored mcc tree plot with gtr on the left and gtri on the right.

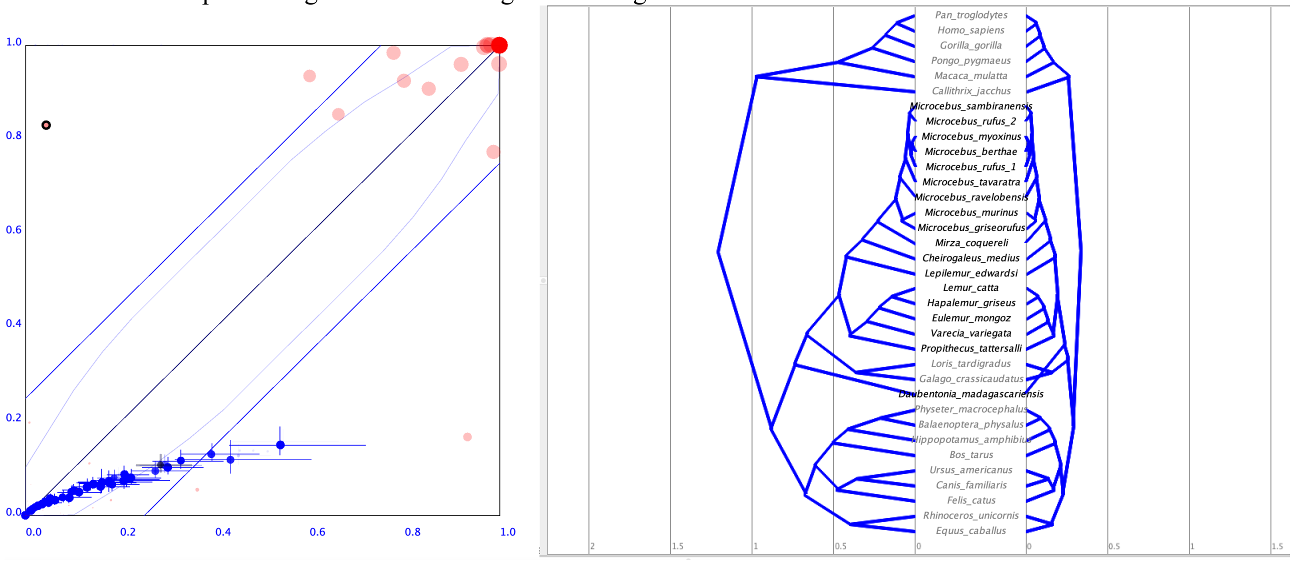


Figure S1.5. (a) A pairwise plot of clade support values (pink) and divergence time estimates (blue crosses). The order is switched here as DensiTree scales its plots off the tree on the left alone. The higher likelihood model (gtr4g) appears on the *x*-axis, and the next lower likelihood model (gtri) appears on the *y*-axis. (b) A mirrored mcc tree plot with gtrg on the left and gtri on the right.

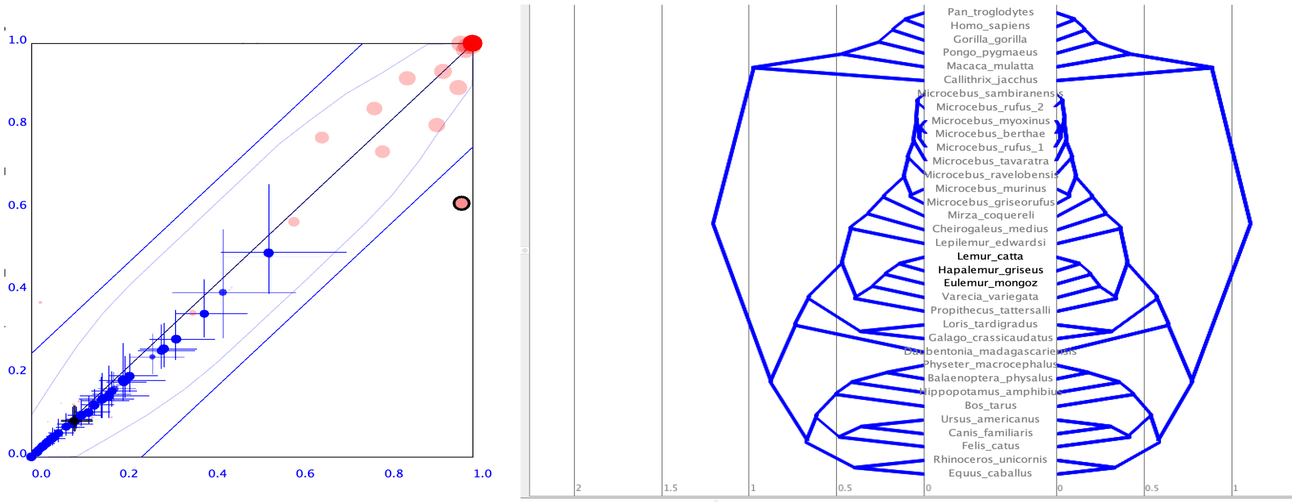


Figure S1.6. (a) A pairwise plot of clade support values (pink) and divergence time estimates (blue crosses). The lower likelihood model (gtr4g) appears on the *x*-axis, and the next lower likelihood model (gtr4gi) appears on the *y*-axis. (b) A mirrored mcc tree plot with gtr4g on the left and gtr4gi on the right. For the final comparison, see figure 3 of the main text.
